# Phylogenomic analyses of Alismatales shed light into adaptations to aquatic environments

**DOI:** 10.1101/2021.11.17.467373

**Authors:** Ling-Yun Chen, Bei Lu, Diego F. Morales-Briones, Michael L. Moody, Fan Liu, Guang-Wan Hu, Chien-Hsun Huang, Jin-Ming Chen, Qing-Feng Wang

## Abstract

Land plants first evolved from freshwater algae, and flowering plants returned to water as early as the Cretaceous and multiple times subsequently. Alismatales is the largest clade of aquatic angiosperms including all marine angiosperms, as well as terrestrial plants. We used Alismatales to explore plant adaptation to aquatic environments by analyzing a data set that included 95 samples (89 Alismatales species) covering four genomes and 91 transcriptomes (59 generated in this study). To provide a basis for investigating adaptation, we assessed phylogenetic conflict and whole-genome duplication (WGD) events in Alismatales. We recovered a relationship for the three main clades in Alismatales as (Tofieldiaceae, Araceae) *+* core Alismatids. We also found phylogenetic conflict among the backbone of the three main clades that could be explained by incomplete lineage sorting and introgression. Overall, we identified 18 putative WGD events across Alismatales. One of them occurred at the most recent common ancestor of core Alismatids, and three occurred at seagrass lineages. We also found that lineage and life-form were both important for different evolutionary patterns for the genes related to freshwater and marine adaptation. For example, several light or ethylene-related genes were lost in the seagrass Zosteraceae, but are present in other seagrasses and freshwater species. Stomata-related genes were lost in both submersed freshwater species and seagrasses. Nicotianamine synthase genes, which are important in iron intake, expanded in both submersed freshwater species and seagrasses. Our results advance the understanding of the adaptation to aquatic environments and whole-genome duplications using phylogenomics.

## Introduction

Land plants evolved from freshwater algae about half a billion years ago (Tena 2020). To adapt to terrestrial environments, land plants acquired genes dealing with intense ultraviolet light, red light, and water evaporation (Han et al. 2019; Kong et al. 2020). Some extant members of major vascular plant lineages have returned to the water as obligates: ferns and lycophytes (e.g. *Azolla*, lsoetaceae), gymnosperms (e.g. *Retrophyllum minus*) and angiosperms (e.g. *Utricularia, Ceratophyllum*, and water lily). The earliest unequivocal remains of angiosperms date to the Early Cretaceous (Friis et al. 2010) and some of the earliest angiosperms are fully aquatic. For example, Nymphaeales have been identified from 125–115 million years ago (mya; Friis et al. 2010) and Ceratophyllales from the late Albian (c. 113 mya; Wang et al. 2018). Aquatic angiosperms constitute only 1–2% of extant angiosperms, but are found in approximately 17% of families representing about 100 evolutionarily independent origins (Cook, 1990). Compared with terrestrial angiosperms, aquatic angiosperms occupy distinctive and stressful ecological habitats characterized by low light levels, reduced carbon and oxygen availability, and mechanical damage through wave exposure. Aquatic angiosperms have adapted to the requirements of aquatic habitats with life-forms notably different from terrestrial plants, generally being divided into emergent, floating-leaved, or submersed forms, the latter most extreme (Les et al. 1997). Survival in aquatic habitats has resulted in adaptations specific to low oxygen concentrations. For example, the aerenchyma in roots, stems, and leaves improves the capture and transportation of oxygen (Jung et al. 2008; Nowak et al. 2010). At the metabolic level, aquatic plants might have enhanced glycolytic fluxes and ethanolic fermentation (Billet et al. 2018).

Aquatic angiosperms comprise freshwater and marine species (often called seagrasses), the latter evolved from the former (Les et al. 1997). A few studies have investigated plant adaptation to freshwater at the genomic level. For example, An et al. (2019) found that disease-resistant genes, which protect plants from pathogens, are not only amplified but also highly expressed in *Spirodela polyrhiza*. Compared with freshwater species, marine species live in environments that have a higher concentration of salt but a lower concentration of iron (Duarte et al. 1995; Anton et al. 2018). Several studies have investigated the genes related to the adaptation from freshwater to seawater. By using genomic data, Golicz et al. (2015), Olsen et al. (2016), and Lee et al. (2016) found that seagrasses lost genes associated with ethylene, terpenoid biosynthesis, ultraviolet damage, and stomatal differentiation. Seagrasses have a constitutive formation of aerenchyma, possibly obviating the need for ethylene production, which is a ubiquitous response to inundation in terrestrial plants to form aerenchyma (Golicz et al. 2015). On the other hand, seagrasses regained genes related to salinity adaptation (Olsen et al. 2016; Lee et al. 2018). However, these studies compared genomic data of seagrasses with those of terrestrial or floating aquatic plants, but not freshwater submersed plants. As the environments for seagrasses and submersed freshwater plants are somewhat similar, a direct comparison between these life forms in the two environments may provide a clearer understanding of loss or gain of genes in aquatic angiosperms and identify genes that are important for adaptation to saline environments. In addition, only a few seagrasses and freshwater species were used in previous studies and the common or unique strategies of aquatic plant adaptation have not been investigated in a more comprehensive and phylogenomic context.

Alismatales, with a worldwide distribution, is the second oldest diverging clade of monocots (Eguchi and Tamura, 2016). It consists of core Alismatids, Araceae, and Tofieldiaceae. Among them, core Alismatids includes 12 families (Alismataceae, Butomaceae, Hydrocharitaceae, Scheuchzeriaceae, Aponogetonaceae, Juncaginaceae, Maundiaceae, Zosteraceae, Potamogetonaceae, Posidoniaceae, Ruppiaceae, and Cymodoceaceae), 58 genera, and c. 593 species (The Plant List: http://www.theplantlist.org/; The Angiosperm Phylogeny Group 2016). Araceae includes 117 genera and c. 3,368 species and Tofieldiaceae includes four genera and 29 species (The Plant List). Alismatales comprises the largest clade of aquatic angiosperms representing all aquatic lifeforms: emergent (e.g. *Alisma*), floating-leaved (e.g. *Spirodela*), and submersed (e.g. *Potamogeton, Najas*). It also encompasses all the marine angiosperms (Zosteraceae, Posidoniaceae, Hydrocharitaceae, and Cymodoceaceae), which comprise three lineages with independent origins from submersed freshwater life-forms (Les et al. 1997). Araceae and Tofieldiaceae also contain a combination of both terrestrial and aquatic taxa. All this makes Alismatales the ideal group for studying aquatic plant adaptations.

Although several studies have investigated the phylogeny of Alismatales, phylogenetic uncertainty remains. One of the recalcitrant relationships is among the three main clades within Alismatales. Analyses using chloroplast (cp) and mitochondrial markers supported a relationship of (Tofieldiaceae, (Araceae, core Alismatids)) (Iles et al. 2013; Ross et al. 2016; Petersen et al. 2015; Luo et al. 2016; Li et al. 2019). However, studies using cp and/or nuclear markers supported a relationship of (Araceae, (Tofieldiaceae, core Alismatids)) (Nauheimer et al. 2012; Hertweck et al. 2015; Azuma and Tobe, 2011; Li et al. 2021). Conflicting phylogenetic relationships also exist in Alismataceae, Potamogetonaceae, and Hydrocharitaceae (Tanaka et al. 1997; Les et al. 2006; Chen et al. 2012; Ross et al. 2016). Investigating the evolutionary history of Alismatales is essential to assess adaptations to aquatic environments.

Whole-genome duplication (WGD; also known as polyploidy) has had a profound influence on plant radiations and environmental adaptations (Van de Peer et al. 2017), such as plant function under salinity (Chao et al. 2013; Yang et al. 2014). WGD events have been identified in two Alismatales lineages: (1) *Zostera* (Zosteraceae), which underwent a WGD around 65 mya (Olsen et al. 2016); and (2) in Araceae, which had WGDs at *Spirodela* and *Pinellia* separately or at their ancestor (Wang et al. 2014; Ren et al. 2018; Qiao et al. 2019). Only *Zostera, Alisma*, and Araceae have been used to evaluate WGDs in Alismatales and the ages of the events discovered are estimated to be older than the stem nodes of the respective lineages where WGD was identified, (e.g. the WGD of *Zostera* (65 mya is older than the stem node age of Zosteraceae (58 mya; Hertweck et al. 2015)). The age inconsistency implies that positions of the WGDs need further assessment. Up to now, studies have not comprehensively explored WGDs within Alismatales and it is still unclear if WGD events are involved in the adaptation to aquatic environments.

Here, we used a data set that included 91 transcriptomes (59 generated in this study) and four genomes. The data set represents 13 of the 14 families within Alismatales (The Angiosperm Phylogeny Group 2016), 40 of the 57 genera within core Alismatids, 17 of the 117 genera within Araceae, and three of the four genera within Tofieldiaceae (The Plant List). The data set also represents terrestrial, freshwater, and marine species. By conducting orthology inference, phylogenetic analyses, and genome duplication inference, we aimed to 1) reappraise the phylogenetic conflict in Alismatales, 2) examine WGD events in Alismatales, especially those potentially involved in adaptation to aquatic environments, and 3) investigate genes associated with adaptation to aquatic and high salinity environments.

## Results and Discussion

A workflow showing data sources, analytical steps, and results is provided in supplementary fig. S1.

### Data Processing and Phylogenetic Analyses

Reads of 59 newly generated transcriptomes were deposited at the NCBI Sequence Read Archive (BioProject: PRJNA809041; supplementary table S1). The number of filtered read pairs ranged from 4.9– 31.5 million. The final number of ‘monophyletic outgroup’ (MO) orthologs with 95 taxa was 1,005. Fifty-three cp genes for 92 samples were also assembled.

RAxML (concatenation) and ASTRAL (coalescent-based) phylogenetic analyses were applied to both the nuclear and cp data sets. The topologies inferred using RAxML (fig. 1) and ASTRAL (fig. 2) with nuclear orthologs are similar except for the position of a *Potamogeton* species. Both analyses resulted in strong support (Bootstrap [BS] =100 or ASTRAL local posterior probability [LPP] =1) for most clades. Both analyses recovered a relationship (Tofieldiaceae, (Araceae, core Alismatids)), however, this lacks BS support (BS =0 and LPP =0.99; fig. 1). There is also low LPP support for one clade within *Potamogeton* (fig. 1).

**Fig. 1.**
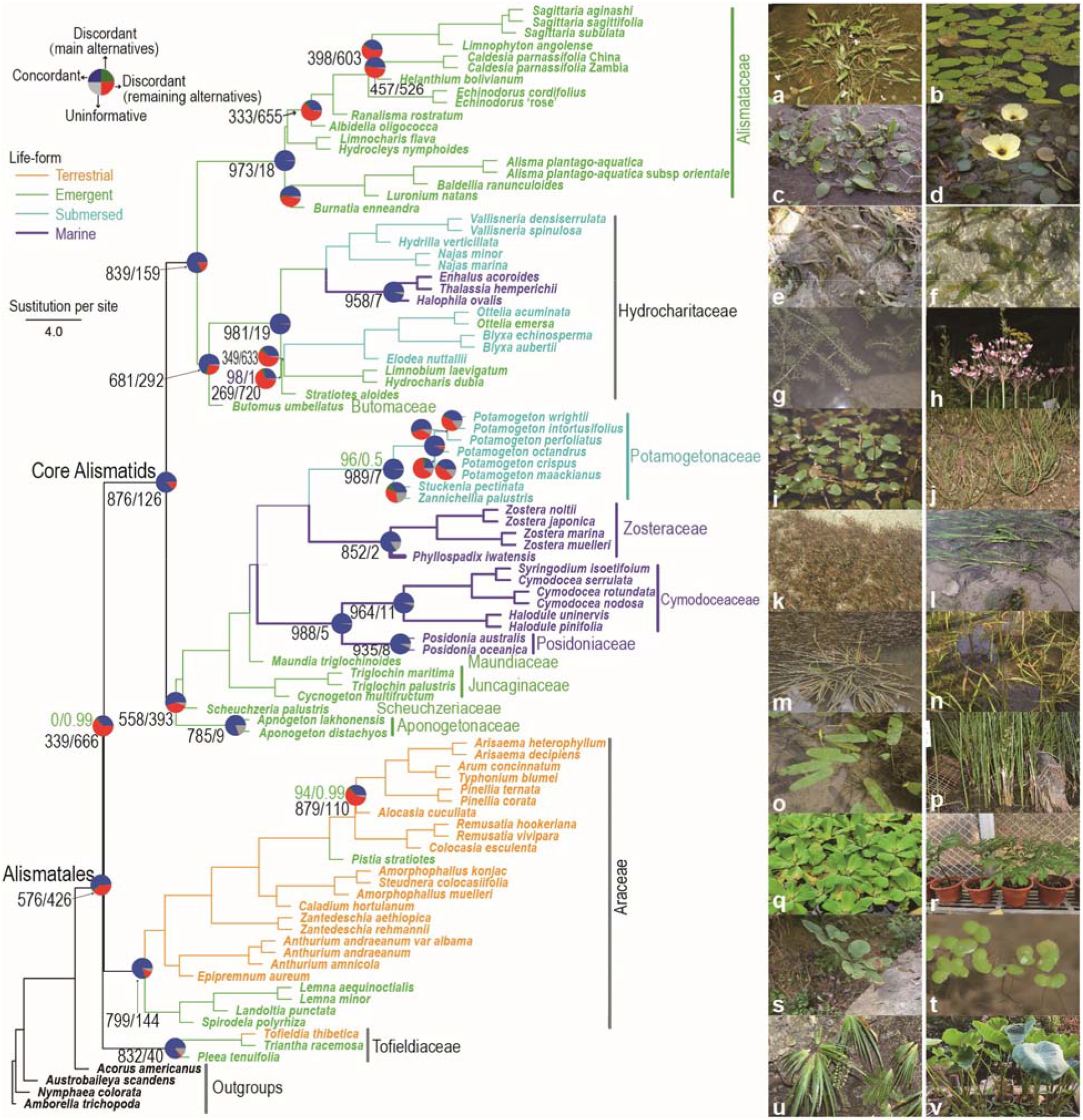
Maximum likelihood phylogeny of Alismatales inferred with RAxML from the concatenated 1,005-nuclear gene matrix. The phylogeny is congruent with the topology inferred with ASTRAL (fig. 2) except *Potamogeton octandrus*. Branches and species names were colored according to the corresponding life-forms. All nodes have maximum support (RAxML bootstrap=100/ASTRAL local posterior probability =1) unless noted near the branches (green). Pie charts represent gene trees for concordant bipartitions (blue), the most frequent alternative topology (green), remaining alternatives (red), and those uninformative for nodes. Numbers below branches represent the number of concordant/discordant genes trees for some nodes. Pie charts and numbers were shown only for nodes discussed in the main text. Plant photos were taken by Lingyun Chen, Jinming Chen, Kuo Liao and Qingfeng Wang. (a) *Sagittaria subulata*, (b) *Albidella oligococca*, (c) *Luronium natans*, (d) *Hydrocleys nymphoides*, (e) *Enhalus acoroides*, (f) *Thalassia hemprichii*, (g) *Hydrilla verticillata*, (h) *Butomus umbellatus*, (i) *Potamogeton distinctus*, (j) *Triglochin maritima*, (k) *Cymodocea rotundata*, (l) *Zostera marina*, (m) *Cycnogeton multifructum*, (n) *Maundia triglochinoides*, (o) *Aponogeton lakhonensis*, (p) *Scheuchzeria palustris*, (q) *Pistia stratiotes*, (r) *Amorphophallus konjac*, (s) *Arisaema lobatum*, (t) *Lemna minor*, (u) *Tofieldia thibetica*, (v) *Colocasia esculenta*. High-resolution plant photos are available at the figshare: https://doi.org/10.6084/m9.figshare.16967767.

**Fig. 2.**
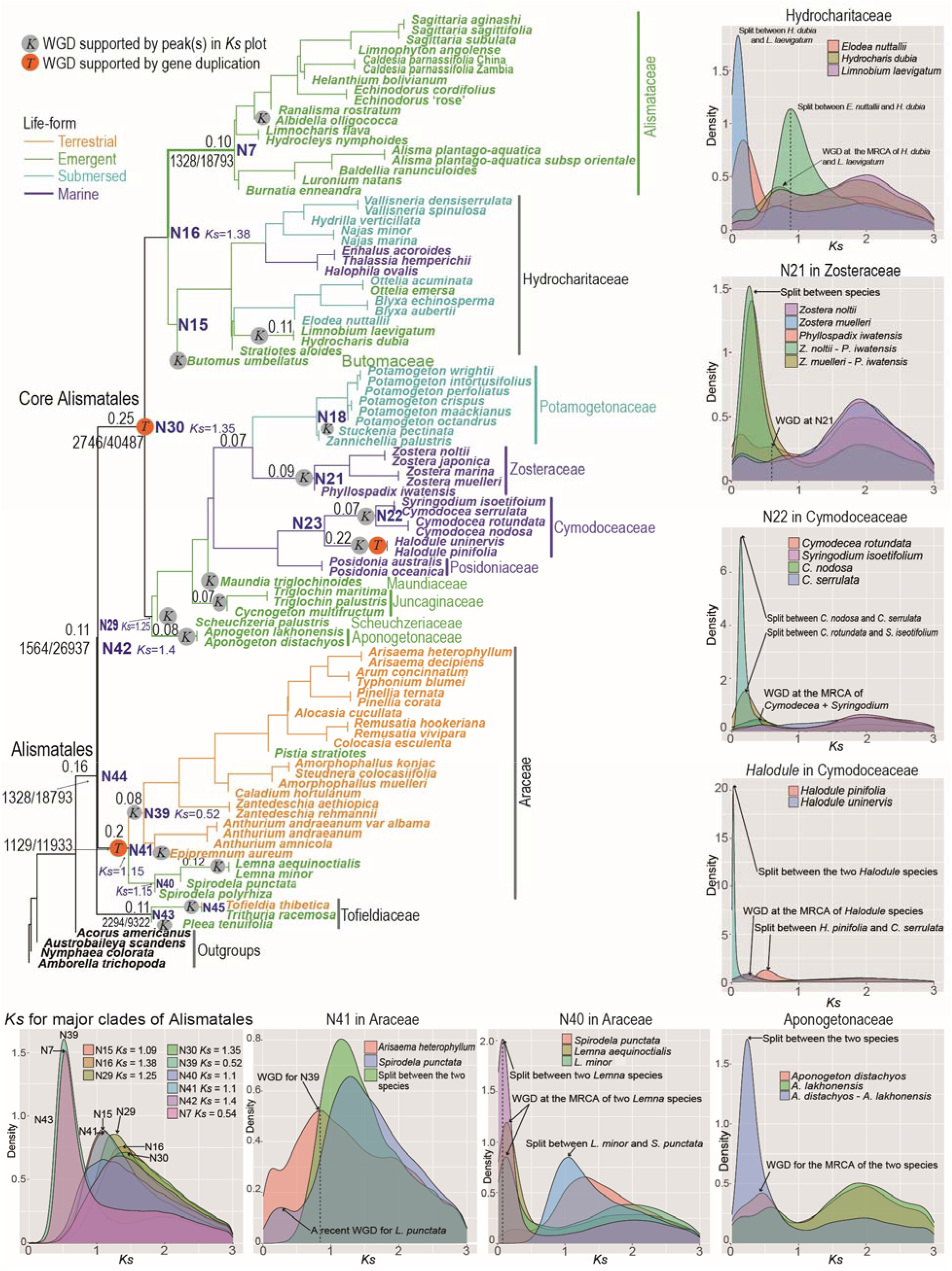
ASTRAL species tree inferred from 1,005 nuclear genes. Percentages next to nodes denote the proportion of duplicated genes (shown when it is ≥7%) when mapping gene clusters to the ASTRAL species tree. Numbers before and behind the virgule (/) represent the number of gene duplications shared by the taxa and the total ancestral gene number before a whole-genome duplication (WGD) event experienced along the branch as inferred from software Tree2GD. The number of gene duplications is shown only when ≥1000. Whole-genome duplications inferred from *Ks* distribution are labeled on the tree. The density plots of within-species *Ks* and between-species *Ks* (inferred using scripts from Yang et al. (2015) and Wang et al. (2019)) are shown for seven representative clades. Node numbers correspond to those in supplementary fig. S6. *Ks* distribution for all species of Alismatales is present in supplementary fig. S8. ‘MRCA’ means most recent common ancestor.

The overall support inferred from cp data is lower than that inferred from the nuclear data set, although most clades have BS >90 or LPP >0.90 (supplementary fig. S2-1 & 2-2). The topologies inferred from nuclear orthologs and cp data are similar. All analyses supported (Tofieldiaceae, (Araceae, core Alismatids)). Deep-level phylogenetic inconsistencies between the nuclear orthologs and cp data exist for *Echinodorus* within Alismataceae and (*Hydrocharis, Limnobium*) within Hydrocharitaceae. Species-level phylogenetic inconsistency was also common within Potamogetonaceae. Almost all the nodes with inconsistencies have BS <90 or LPP <0.90 (supplementary fig. S2-1 & 2-2).

We combined cp coding sequences (CDSs) assembled from RNA sequencing (RNA-seq) reads and CDSs extracted from chloroplast genomes for phylogenetic analyses. The topology of our cp trees (supplementary fig. S2-1 & 2-2) was similar to that inferred from chloroplast genomes (Ross et al. 2016). For example, (*Hydrocharis* + *Limnobium*) was clustered with a clade formed by *Thalassia, Enhalus, Halophila, Vallisneria*, and *Najas*. Since many plant plastomes are heavily RNA-edited and a combination of DNA sequences with RNA-edited transcripts may cause problems in phylogenetic analyses (Ramaswami et al. 2013), the phylogeny using plastome CDSs potentially result in bias or error of branch length. Therefore, we used the nuclear phylogeny for further analyses in our study.

Overall, the phylogenetic analyses supported monophyly for all families of Alismatales for which multiple accessions were sampled (i.e. Alismataceae, Aponogetonaceae, and Araceae) and the Core Alismatids, consistent with previous studies (Les et al. 1997; Iles et al. 2013; Ross et al. 2016; Petersen et al. 2015; Luo et al. 2016).

### Phylogenetic Conflict

To investigate phylogenetic conflict, we calculated the concordant and conflicting bipartitions, and internode certainty all (ICA; Salichos et al. 2014) values using PhyParts. ICA value quantifies the degree of conflict in a given internode. An ICA value closer to 1 indicates strong concordance in the bipartition of interest, while an ICA value closer to 0 indicates equal support for one or more conflicting bipartitions. A negative value indicates that the internode of interest conflicts with one or more bipartitions that have a higher frequency, and an ICA value close to -1 indicates the absence of concordance for the bipartition of interest (Salichos et al. 2014). We also calculated Quartet Concordance (QC), Quartet Differential (QD), and Quartet Informativeness (QI) scores using Quartet Sampling (QS; Pease et al. 2018). A QC value closer to 1 indicates all concordant quartets, a QC closer to 0 indicates equivocal concordant/discordant quartets, and a QC smaller than 0 indicates discordant quartets are more frequent than concordant quartets. A QD of ‘-’ indicates no alternative topologies (i.e. QC = 1), a QD closer to 1 indicates two alternative histories appear equally frequently, and a QD closer to 0 indicates one of the two alternative histories is favored. A QI value close to 1 indicates all quartets are informative, while a value close to 0 indicates all quartets were uncertain (Pease et al. 2018).

The monophyly for most families of Alismatales was highly supported with at least 80% of informative gene trees being concordant (fig. 1), ICA >0.8, and full QS support (1/-/1) (supplementary fig. S3). Conflict was detected for the clade formed by core Alismatids and Araceae (number of concordant/conflicting bipartitions is 339/666; ICA =0.3; QS score =-0.24/0.14/0.99). Conflict was also detected for another few clades, for example, the two sub-clades within Alismataceae and the clades involving *Stratiotes* (ICA =0.43, QS =0.042/0/1) and *Limnobium* + *Hydrocharis* (ICA =0.5, QS = - 0.14/0.67/0.99) within Hydrocharitaceae (supplementary fig. S3).

To detect possible reticulation on the nodes with large phylogenetic conflict, we performed species network searches using PhyloNet v3.8.2 (Than et al. 2008) with a Bayesian Markov chain Monte Carlo (MCMC) algorithm for Alismatales, Alismataceae, Hydrocharitaceae, Potamogetonaceae, and Araceae. All five independent PhyloNet runs for Alismatales, Hydrocharitaceae, and Potamogetonaceae converged, while four of 10 runs for Alismataceae and five of 10 runs for Araceae converged (supplementary fig. S4). The MPP (maximum posterior probability) networks were obtained after summarization of converged runs (fig. 3). PhyloNet calculates inheritance probabilities (γ), by which γ ≈0.5 infers hybridization, γ ≈0.1 infers introgression, and a small γ (<0.05) infers incomplete lineage sorting (ILS) (Solís-Lemus et al. 2017). We discuss each of the five lineages below.

**Fig. 3.**
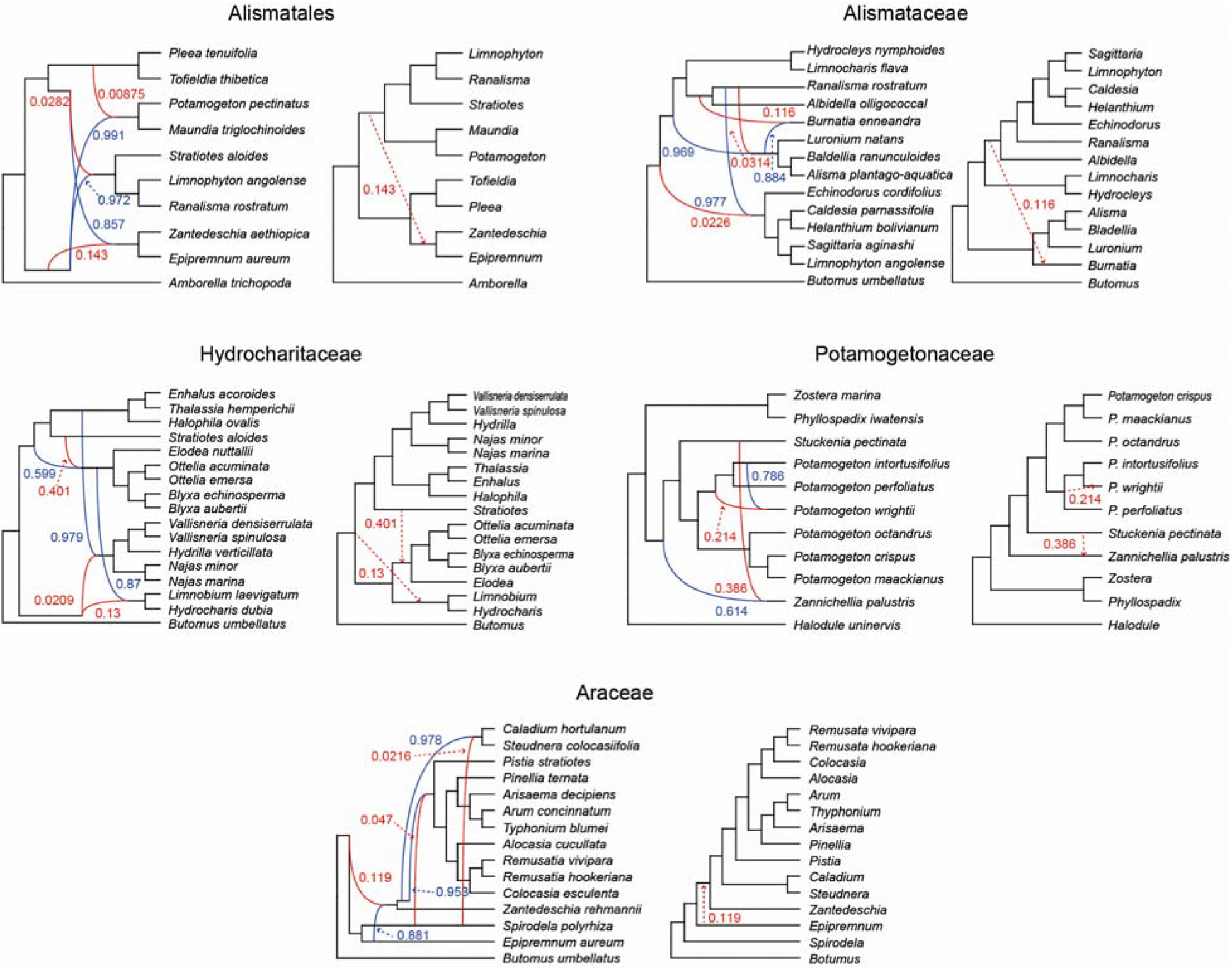
Maximum posterior probability species networks of the five reduced-taxon data set inferred with PhyloNet. Numbers next to curved lines indicate inheritance probabilities (γ) for each hybrid node. Red (γ<0.5) and blue (γ >0.5) curved lines indicate the minor and major hybrid edges respectively. Trees next to each network represent the networks showing only reticulation events with a minor edge with inheritance probability ≥0.1.

#### Alismatales

The MPP topology from PhyloNet recovered the relationship (core Alismatids, (Araceae, Tofieldiaceae)) (H1; fig. 3). The AU tests do not show support for any of the three possible tree topologies for Alismatales as 626 of the 798 were equivocal (fig. 4a). However, H1 was more frequent when counting the number of trees supporting each of the three possible topologies by significant likelihood support (fig. 4b). Moreover, H1 was also more frequent when counting the number of trees using BS values (fig. 4c). The other two topologies for Alismatales were not supported: 1) The topology of hypothesis H0 (Tofieldiaceae, (Araceae, core Alismatids)) was recovered in phylogenetic analyses using both nuclear and cp data (fig. 1, supplementary fig. S2). However, the clade Araceae + core Alismatids lacked BS support (fig. 1) and had low ICA and QS scores (supplementary fig. S3). PhyloNet found that Araceae inherited about 14.3% (γ =0.143; fig. 3) of its genome from the most recent common ancestor (MRCA) of core Alismatids. Given the γ value, introgression explains the close relationship between Araceae and core Alismatids (H0) in our results and previous studies (Iles et al. 2013; Ross et al. 2015; Petersen et al. 2015; Luo et al. 2016). 2) The hypothesis H2, (Araceae, (Tofieldiaceae, core Alismatids)), was recovered but weakly supported in previous studies. Namely, the nodal support for Tofieldiaceae + core Alismatids was low, BS <75 in Azuma and Tobe (2011), Nauheimer et al. (2012), and Liang et al. (2019). The PhyloNet analyses found that core Alismatids inherited 0.875% (γ =0.00875) of its genome from Tofieldiaceae and 2.82% (γ =0.0282) of its genome from the MRCA of Tofieldiaceae + core Alismatids. Given the low γ values, ILS best explains the close relationship between Tofieldiaceae and core Alismatids (H2). 3) A polytomy hypothesis for Alismatales was rejected, as polytomy tests recovered all nodes have p-values =0. Given the low nodal support and short branch length for the clade core Alismatids + Araceae and AU test results, the evolution of these three clades of Alismatales may be the result of a rapid radiation, which makes resolution of the topology difficult. In conclusion, our AU analyses slightly favored topology H1 for Alismatales, but there is phylogenetic conflict regarding the three major clades of Alismatales that could be due to introgression and ILS.

**Fig. 4.**
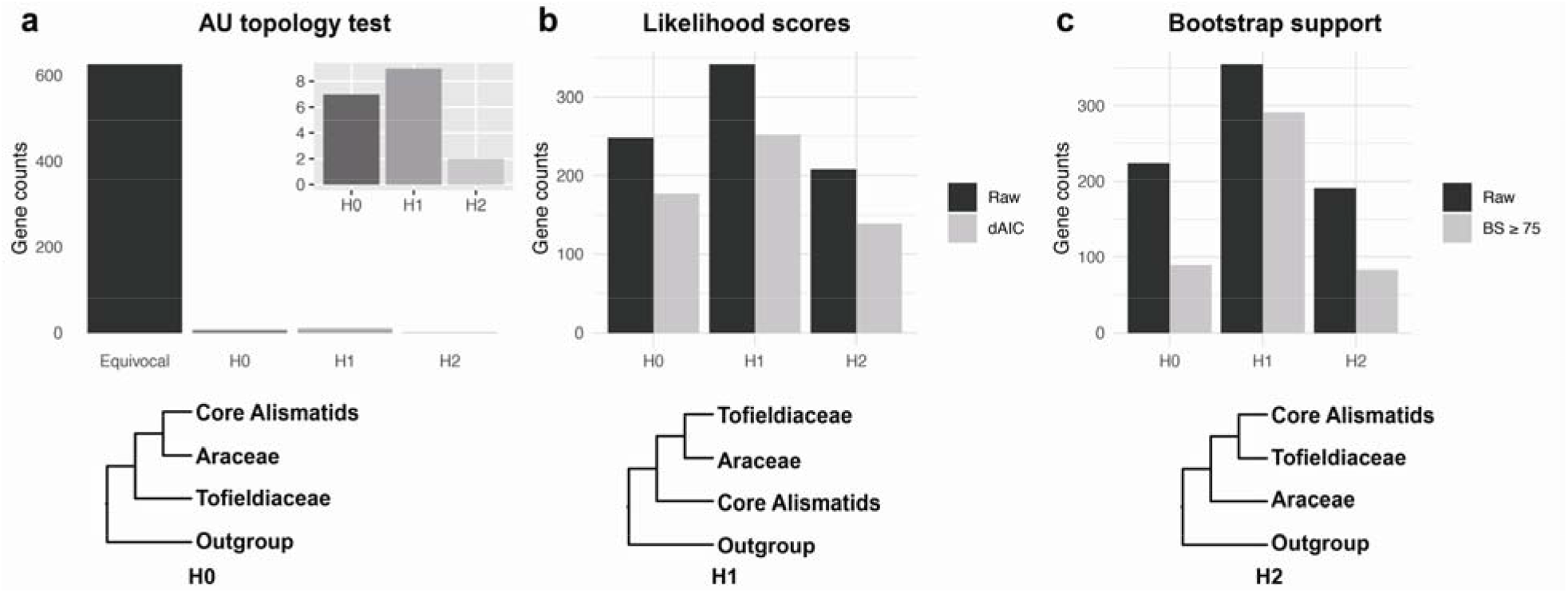
Topology test of the three main clades within Alismatales. Gene counts for topology tests of the three main clades using the Approximately Unbiased (AU) test (a), RAxML likelihood scores (b) and bootstrap support values (c).

#### Alismataceae

The major phylogenetic inconsistency within Alismataceae is the position of *Burnatia*. Using three cp markers, *Burnatia* was resolved as a sister to *Hydrocleys* + *Limnocharis* (Chen et al. 2012). However, using additional cp markers, *Burnatia* was recovered as sister to a clade comprising *Alisma, Baldellia, Damasonium* and *Luronium* (Lehtonen et al. 2017; Li et al. 2022), consistent with our analyses (fig. 1 & 2). PhyloNet inferred three possible reticulations in Alismataceae (fig. 3). Two of them have minor edges with small γ-values (0.0226 and 0.0314, fig. 3), which supports ILS. The third reticulation recovered that *Burnatia* inherited about 11.6% (γ =0.116) of its genome from the MRCA ancestor of *Ranalisma* + *Albidella*. Moreover, *Ranalisma* + *Albidella* was phylogenetically close to *Hydrocleys* + *Limnocharis* (fig. 1). The close relationship between *Burnatia* and *Hydrocleys* + *Limnocharis* (Chen et al. 2012) could be due to introgression from the MRCA of *Ranalisma* + *Albidella* to *Burnatia*.

#### Hydrocharitaceae

Relationships among the major clades of Hydrocharitaceae have been inconsistent across multiple studies. The family comprised two major clades when only cp markers were used, one included *Halophila, Thalassia, Enhalus, Najas, Hydrilla*, and *Vallisneria*, another included *Egeria, Elodea, Blyxa, Ottelia, Lagarosiphon, Stratiotes*, and *Limnobium* + *Hydrocharis* (Tanaka et al. 1997). However, *Limnobium* + *Hydrocharis* was clustered with the former clade by Chen et al. (2012), contradicting Tanaka et al. (1997). Les et al. (1997; 2006) resolved *Limnobium + Hydrocharis* to be sister to all other Hydrocharitaceae, but with poor support. In our ASTRAL analyses, *Limnobium + Hydrocharis* had conflict (ICA=0.5, QS=0.14/0.67/0.99) (supplementary fig. S2). The PhyloNet analysis recovered one reticulation with γ =0.13, suggesting introgression from the MRCA of Hydrocharitaceae to the *Limnobium + Hydrocharis* ancestor. Thus, introgression possibly led to the phylogenetic inconsistency in the placement of *Limnobium + Hydrocharis. Stratiotes* was resolved as a sister to all other Hydrocharitaceae taxa (Chen et al. 2012; Ross et al. 2016; Li et al. 2020), contradicting Tanaka et al. (1997), which clustered it with the clade formed by *Egeria, Elodea, Blyxa, Ottelia*, and *Lagarosiphon*. In our analyses, the clade including *Stratiotes* also showed conflict (ICA=0.43, QS=0.042/0/1). The PhyloNet analysis recovered a hybridization event (γ =0.599 and 0.401) between *Stratiotes* and the MRCA of the clade comprising *Elodea, Ottelia*, and *Blyxa*, which can explain the phylogenetic inconsistency of *Stratiotes*. Overall, phylogenetic conflict regarding *Limnobium + Hydrocharis* and *Stratiotes* can be explained by introgression and hybridization, respectively.

#### Potamogetonaceae

This is a taxonomically complex group with several hybrid taxa reported. Based on internal transcribed spacer (ITS) and cp data, *Potamogeton wrightii* was reported as a parent species of multiple species, including *P. biwaensis* (Iida et al. 2018) and *P. anguillanus* (Du et al. 2010). Our PhyParts and Quartet Sampling analyses suggested strong phylogenetic conflict within Potamogetonaceae (high proportion of discordant bipartitions, fig. 1; low ICA and QS scores, Supplementary fig. S3). The PhyloNet analyses found that *P. wrightii* is a hybrid between *P. intortusifolius* and an extinct or unsampled taxon with γ 0.214 (fig. 3). Moreover, *Zannichellia palustris* was recovered as a potential hybrid between *Stuckenia pectinata* and an extinct or unsampled taxon (γ = 0.386). The phylogenetic conflict could be attributed to hybridization, but the extent of hybridization will require more comprehensive sampling.

#### Araceae

Introgression was detected (fig. 3) from an extinct/unsampled Araceae ancestor to the clade that included all of the sampled taxa except *Epipremnum* and *Spirodela* (γ =0.119). Based on cp markers and a few nuclear markers, introgression has been detected for Araceae, such as *Cryptocoryne* species (Jacobsen et al. 2019). These results suggest that introgression affected the relationships at the genus-level, which has not previously been reported in Araceae.

### Divergence time estimation

To get a timescale for Alismatales diversification, we performed a BEAST analysis using eight fossils and one secondary calibration (supplementary table S2). Divergence time estimation dated the crown age of Alismatales around 135.7 mya (95% HPD: 119.4–153.9; supplementary fig. S5). This age is slightly older than that in recent studies, such as 124 mya in Givnish et al. (2018) and 122 mya in Hertweck et al. (2015), but is younger than that of Li et al. (2019) at around 148 mya. The stem node age of the seagrass clade within Hydrocharitaceae was around 62.8 mya. The stem node ages of the seagrass lineages Zosteraceae and Cymodoceaceae + Posidoniaceae were at 51.7 mya (43.0–60.7) and 67.3 mya (56.8– 78.4), respectively. These ages are consistent with previous studies. For example, divergence time estimation with fossil calibrations found the mean stem node age of Zosteraceae at 53.5 mya (Hertweck et al. 2015). However, these estimated ages are younger than the oldest seagrass fossils reported from the Late Cretaceous (Larkum et al. 2018), which could be attributed to the view that some seagrasses have gone extinct over evolutionary time (Larkum et al. 2018).

### Whole genome-duplications are common across Alismatales

To infer WGDs, we first calculated the proportion and number of duplicated genes for each node in the species tree using the tree-based methods (script *map_dups_mrca*.*py*) of Yang et al. (2018) and Tree2GD. A high proportion of duplicated genes at a node could be due to a WGD at that node (type I duplication; Huang et al. 2016). However, a WGD that happened at one subclade within a node, but lacking from another, could also lead to the detection of a high proportion of gene duplication at that node (type II or type III duplication; supplementary table S3; Huang et al. 2016). Compared with the type II and type III duplication, type I is more likely a real WGD (Huang et al. 2016). Tree2GD calculated the proportion of genes supporting each of the three types. For tree-based analyses, we used ≥20% duplicated genes (Yang et al. 2018) as a cutoff to identify WGD. Type of duplication (Huang et al. 2016) and evidence from the literature were also considered.

Results showed that three clades have ≥20% duplicated genes: core Alismatids (N30, 25%/2746), Araceae (N41, 20%/1129), and *Halodule* (Cymodoceaceae, 22%/NA; fig. 2 and supplementary fig. S6). Core Alismatids had a higher proportion of type I duplications than the other two types (supplementary table S3), which implies a WGD could have occurred (supplementary table S3) at that node. Araceae has a higher proportion of type II or type III duplication than type I. To verify the candidate WGD event at the MRCA of core Alismatids, Zosteraceae, and Araceae respectively, we reconstructed a phylogeny using 99 samples, then mapped 3,493 gene clusters to the phylogeny and counted the number of duplicated gene clusters at each node of Alismatales using PhyPart (Smith et al. 2015). The results showed that 663 out of 3,493 gene clusters were duplicated at the MRCA of the core Alismatids (supplementary fig. S7). Among the 663 gene clusters, 148 (22.3%) gene pairs have syntenic evidence in *Z. marina*. For the WGD at Zosteraceae (N21), 205 duplicated gene clusters were found, and 128 (62.4%) gene pairs have syntenic evidence in *Z. marina*. For the WGD at Araceae (N41), 552 duplicated gene clusters were found, and 203 (36.8%) gene pairs have syntenic evidence in *S. intermedia* (supplementary fig. S7). The WGD at the MRCA of Araceae was supported by Wang et al. (2014) and An et al. (2019), which suggested that two rounds of WGDs in *Spirodela* occurred at c. 95 mya, consistent with the age of Araceae (mean stem node age: 128.0 mya; mean crown node age: 82.3 mya). A WGD could have occurred at the stem node of *Halodule*, which had 22% duplicated genes, although the type of WGD could not be calculated given only two taxa in this clade. We propose that there are three candidate WGD events, (1) the MRCA of core Alismatids, (2) the MRCA of Araceae, and (3) the MRCA of *Halodule*.

As an alternative method to infer WGDs, we also plotted the distribution of synonymous substitutions (*Ks*). We consider a *Ks* peak as a candidate WGD if it is clear in within-species *Ks* plot and between-species *Ks* plot. For an obscure peak, evidence from the literature is required. The analyses suggested 16 candidate WGD events (fig. 2, supplementary fig. S8). No *Ks* peak was identified for the candidate WGD at core Alismatids (N30) and Araceae (N41). The failure to detect the WGD at these nodes could be due to saturation effects in *Ks* estimation (Rabier et al. 2014). We discuss the major candidate WGDs identified based on *Ks* distribution below.

In total, 16 WGD events were identified using the *Ks* method. A *Ks* peak around 0.7 (supplementary table S4) was recovered for 20 of 21 species at node N39 within Araceae (supplementary fig. S8-3). The peak at 0.7 is younger than the diversification peak of Araceae (*Ks* =1.15), but older than the diversification of N39 (*Ks* =0.52; supplementary table S5). N39 has 8% duplicated genes (fig. 2) and type I accounts for the highest proportion of duplication. Therefore, there should be a WGD for N39. This WGD was supported by a WGD in *Pinellia* (within the N39) identified by Ren et al. (2018). The *Ks* plot recovered a clear peak of 0.15 for *Lemna minor* and *L. aequinoctialis* (Araceae), older than the split between the two species (*Ks* =0.1), but younger than the split between *Lemna* and *Spirodela* (*Ks* =1.1). *Lemna* has 12% duplicated genes (fig. 2). Therefore, a WGD event may have occurred for *Lemna*, consistent with the opinion that at least one WGD occurred for the ancestor of *Lemna* after the split between *Lemna* and *Spirodela* (Van Hoeck et al. 2015). We also detected candidate WGD events for *Albidella olligococcal* (Alismataceae; supplementary fig. S8-1; supplementary table S4), *Limnobium laevigatum* + *Hydrocharis dubia, Butomus umbellatus* (Butomaceae), *Stuckenia pectinata* (Potamogetonaceae), *Maundia triglochinoides* (Maundiaceae), *Cycnogeton multifructum* (Juncaginaceae; supplementary fig. S8-2), *Scheuchzeria palustris* (Scheuchzeriaceae), Aponogetonaceae, and Tofieldiaceae (supplementary fig. S8-3). These WGD events were not detected when using tree-based methods, which may in part be because these methods need more comprehensive sampling.

Among the 16 WGD events identified using *Ks*, some occurred for seagrass lineages, namely Zosteracae and Cymodoceaceae. Olsen et al. (2016) suggested a WGD for *Zostera* (Zosteraceae) occurred around 65 mya. Our analyses recovered a peak (*Ks* ≈0.6) for three of the six species in Zosteraceae. Between-species *Ks* plots showed that the peak is at the node of Zosteraceae (N21, 9% duplicated genes), which has a stem node age of 48–65 mya. Therefore, the WGD at Zosteraceae identified here could correspond to the WGD at *Zostera* reported by Olsen et al. (2016). Cymodoceaceae has two candidate WGD events. One is at the MRCA of N22 (*Syringodium* + *Cymodocea*), supported by a peak (*Ks* ≈0.45) of all the four species and 7% duplicated genes at the node (Fig. S8-2). Another is at *Ks* ≈0.25, positioned at the *Halodule* clade and supported by 22% duplicated genes. Reconstruction of the evolution of chromosome numbers for marine angiosperms suggested that WGD events had occurred in Zosteraceae and Cymodoceae (Silva et al. 2021).

In summary, we identified 18 candidate WGD events (supplementary table S4). Among them, two (N30 and N41) were identified using tree-based methods, 15 were identified using *Ks*, and one (*Halodule*) was identified using both methods. WGD is believed to have evolutionary consequences for adaptation and is well documented for plants (Hollister. 2015) and has been thought to have particular importance for adaptation to novel environments (Van de Peer et al. 2017). Recent genomic studies have provided empirical evidence for WGDs contributing to adaptation and diversification of mangroves that grow in aquatic high saline environments, such as tropical coastal swamps (Xu et al. 2017; Feng et al. 2021) as well as other novel or harsh environments for plants (Feng et al 2020, Wang et al 2019). The frequency of WGDs among the Alimatales provides further evidence that WGDs could have also contributed to novel environments and could have a role in the adaptation to aquatic habitats and given the multiple inferred events among the seagrasses could have a role in the adaptation for this particularly extreme environment as inferred for mangroves (Feng et al. 2021). However, WGD events do not appear to be associated with specific life-forms among the aquatic lineages, as all four life-forms in Alismatales have WGD events. More broad analyses using a phylogenomic context across hydrophyte lineages will provide more insight into the role of WGD in these adaptations.

### Genes involved in aquatic adaptation

Previous studies have reported genes possibly related to the adaptation of aquatic environments (Olsen et al. 2016; Han et al. 2019). The evolutionary patterns of gene loss or gain within different aquatic plants remain poorly known. To investigate the evolutionary patterns of these genes (supplementary table S6), we compared the numbers of gene copies for eight gene groups in Alismatales species under a phylogenetic context in relation to the four life-forms found in Alismatales. These results largely corroborate the results of Olsen et al. (2016) regarding the seagrass *Z. marina*. However, our in-depth phylogenetic sampling of Alismatales, incorporating multiple submersed and marine life-form plants, provides evidence that there is a great deal of variation within and among life-forms regarding gene gain and loss and some genes previously considered to be lost or directly related to adaptation to the aquatic environment should be reconsidered.

1. Ethylene is a critical plant hormone that controls various physiological responses (Broekgaarden et al. 2015). The biosynthesis and reception of ethylene mainly involve gene *ACS, ACO, ETR1, ERS1, ETR2, ERS2*, and *EIN4, CTR1*, and *EIN2*. Olsen et al. (2016) attributed the loss of ethylene-related genes in *Z. marina* to the loss of stomata and the absence of insect herbivores in the sea. Our analyses found that all of the targeted ethylene-related genes were lost in *Z. marina* and *Z. muelleri*, consistent with Olsen et al. (2016), and loss of these genes was generally more common among seagrasses than freshwater taxa with notable exceptions in *Posidonia* and two Hydrocharitaceae genera (Fig. 5b). The only freshwater taxon, *Elodea nuttallii*, with comparable losses is known for tolerance to brackish water (Thouvenot and Thiébaut 2018) (fig. 5b & 5c-i–iv). The loss of these genes is not notable in freshwater taxa in comparison to terrestrial Alismatales, suggesting it is not directly related to stomatal loss as previously suggested (Olsen et al. 2016). There are seagrass pathogens in the sea (Sullivan et al. 2018) and animals feeding on seagrasses, such as snails (Rotini et al. 2018). The ethylene in seagrasses could respond to pathogens and grazers. It is also possible that ethylene plays a role in seagrass growth and development. Overall, complete loss of ethylene-related genes is not found in all seagrasses, but the loss is common across most lineages and more are lost in seagrasses than freshwater taxa (supplementary table S7).
2. Terpenoids play an important role in plant-insect and plant-pathogen interactions, as well as plant development (Keeling and Bohlmann 2012). The terpene synthase (*TPS*) gene family examined here, comprises five sub-families, a, b, c, e/f, and g (Jiang et al. 2019). *TPS* sub-family a and g were lost in all core Alismatids representing an ancestral state for the clade. Compared to terrestrial plants, subfamily b, c, and e/f are more common in the emergent life-form plants (fig. 5b & 5c-v). This follows a similar pattern to the ethylene-related ACS type 2 and ERS sub-clade 2 gene (fig. 5b). Both ethylene and terpenoids are essential for a response to stress. Emergent plants need to be able to adapt to extreme shifts in the environment during their lifetime and be able to regulate metabolic processes under both inundated and emergent conditions. The retention of subfamily b, c, and e/f and more ethylene-related genes in emergent plants could be attributed to the conditions they live in. This result is corroborated by a study in *Ranunculus*, the emergent aquatic plant *R. sceleratus* has more genes under positive selection than its terrestrial and submersed relatives (Zhao et al. 2016).
3. Aerenchyma-related genes. Enhanced Disease Susceptibility (EDS1) and Phytoalexin Deficient 4 (PAD4) are regulators for lysigenous aerenchyma formation by programmed cell death in *Arabidopsis* (Mühlenbock et al. 2008). Regulator of G protein signaling (RGS) is an inducer for aerenchyma formation in the absence of ethylene in maize (Steffens and Sauter 2010). Aerenchyma in aquatic plants is present constitutively in leaves and roots as an adaptation, thus *EDS1, PAD4* and *RGS* may be expected to have an important role in the submersed and marine life-form. Our results showed that *EDS1* was present in 15 of the 21 terrestrial species of Alismatales, but were lost in 68 of the 70 aquatic samples of Alismatales (fig. 5b). *PAD4* was only present in two species of Tofieldiaceae. *RGS* was present in 20 of the 21 terrestrial species of Alismatales, but were lost in 44 of the 70 aquatic samples of the same group (fig. 5b). Interestingly, the Kyoto encyclopedia of genes and genomes (KEGG) orthology (abbreviated as KO) terms involving aerenchyma were enriched in submerged life-forms in our KO analyses. These results imply that the formation of aerenchyma in aquatic plants of Alismatales may be independent of *EDS1, PAD4* and *RGS*, and these genes may be more important for an induced response rather than the constitutive production of aerenchyma and therefore lost.
4. Stomata-related genes. Stomata control transpiration and regulate gas exchange necessary for photosynthesis. *SPCH, MUTE*, and *FAMA* genes (together referred to as *SMF*), belong to the bHLH super-family, control the stomatal differentiation from meristemoid mother cells (MMC) to guard cells (GC) (Pires et al. 2010; Ran et al. 2013). *ICE*/*SCRM* genes, also belong to bHLH; they control stomatal development and are also involved in the cold acclimation response and freezing tolerance (Pires et al. 2010). While stomata are ubiquitous in terrestrial and aquatic plants with emergent or floating leaves, they are absent from submersed leaves. We found that gene fraction for *SMF* genes decreased (or the genes were lost) in the submersed and marine life-form (fig. 5c-viii), which can be attributed to the absence of stomata. *SMF* genes were also absent from some species that have stomata, such as *S. polyrhiza*, a species with whole genomic data accessed in this study. The gene loss can be attributed to the multifunctionality of *SMF* genes, as a single *SMF* gene could control the specification from MMC to GC in early land plants (Ran et al. 2013). Alternatively, the function of *SMF* genes could have been supplemented by the *ICE*/*SCRM* genes (Ran et al. 2013; Olsen et al. 2016), which have a higher gene fraction in the submersed and marine life-forms than in the emergent life-form (fig. 5c-ix).
5. Nicotianamine, a metabolite synthesized in plants, has a critical role under iron and zinc deficiency in plants (Klatte et al. 2009; Schuler et al. 2012). Our results showed that the gene fraction of nicotianamine synthase (NAS) genes increased in submersed plants and seagrasses (fig. 5b & 5c-x). For example, 14 of the 16 marine species sampled have 1–2 gene copies (supplementary table S7), while no copies were found in most terrestrial or emergent life-forms. The iron concentration of seawater is only one percent of that in rivers and groundwater (Firdaus et al. 2014) and iron deficiency was found to be common in Caribbean seagrasses, Red Sea seagrasses, and macroalgae (Duarte et al. 1995; Anton et al. 2018), which restricted their productivity. Although the reason(s) for the increase of NAS copies in submersed plants is unclear, the increase in seagrasses could be attributed to the iron deficiency in seawater.
6. Light-related genes. The light-harvesting chlorophyll a/b-binding (LHCB) protein 1 belongs to the HLCB super-family, which regulates photosynthetic light-harvesting and several non-photochemical processes (Xu et al. 2012). Our analyses suggest that *LHCB1* genes expanded in *Z. marina* as reported by Olsen (2016). We also found *LHCB1* expanded at the MRCA of core Alismatids, however, it contracted among most other marine lineages (fig. 5b & 5c-xi; supplementary table S7). These results do not support the enhancement of these gene families due to low light alone, but provide insight into variation in adaptive strategy for marine lineages. Land plants evolved phytochromes, cryptochromes, and UVR8 to sense and capture light and overcome intense ultraviolet light in the terrestrial condition (Han et al. 2019). Our analyses suggested that *phytochrome C* is absent in most seagrasses (except *Posidonia* and Hydrocharitaceae), whereas *cryptochrome 2* (*CRY-2*) and *UVR-8* are absent in *Zostera*, as reported by Olsen et al. (2016), but present in most other seagrasses (fig. 5b; fig. 5c-xiii & xiv). Olsen et al. (2016) attributed the loss of these genes to the differences in light absorption by plants through water. However, our results suggested that these genes are widely retained in submersed plants and seagrasses, despite their underwater life-form. Further exploring the effects of plant depth and diffusion of light through high salinity water will be important to explore gene function regarding these environments.
7. Cell wall-related genes. Sulfated polysaccharides (SPs) are cell wall components. They were mainly found in algae (McCandless and Craigie, 1979), and lost in land plants (Olsen et al. 2016). The SPs were synthesized by aryl sulfotransferases (AS). Olsen et al. (2016) found that *Z. marina* regained the aryl sulfotransferase genes as an adaptation to the marine environment. We found that all three major clades within Alismatales have regained *AS* genes (fig. 5b; fig. 5c-xv), suggesting these genes could be involved in the adaptation to aquatic environments in general. The AS genes were also found in all seagrass lineages sampled here, corroborating the chemical studies that SPs exist in seagrass Cymodoceaceae, Hydrocharitaceae, Zosteraceae, and Ruppiaceae (Aquino et al. 2011). SPs have also been identified in freshwater plants such as *Eichhornia crassipes, Hydrocotyle bonariensis* and *Nymphaea ampla* (Dantas-Santos et al. 2012). No SPs have been reported from the freshwater plants within the core Alismatids. More chemical studies are needed to detect if these plants have SPs. The cuticle covers aerial surfaces as a plant adaptation to terrestrial habitats (Kong et al. 2020). The biosynthesis of cuticles involves more than 32 genes (Kong et al. 2020). We found there is no significant difference in copy number for these genes across Alismatales except for the *BODYGUARD* (*BDG*) gene. The *BDG* encodes an epidermis-specific cell wall-located protein that is required for the formation of the cuticle in *Arabidopsis* (Kurdyukov et al. 2006). The *BDG* has contracted in the submersed and marine life-forms (fig. 5c-xvi), which can be attributed to the much reduced or absent cuticle found in these plants (Enriquez et al. 2014).
8. Osmoregulation. H^+^-ATPases (AHA) are ion pumps that use the energy of ATP hydrolysis to reduce Na + influx, which plays a major role in salt stress tolerance. Olsen et al. (2016) found the gene encoding AHA expanded in *Z. marina* and our results suggested this is a common strategy in seagrasses, as the gene copy number increased in all the four seagrass clades (supplementary table S7 & fig. 5c-xvii).

**Fig. 5.**
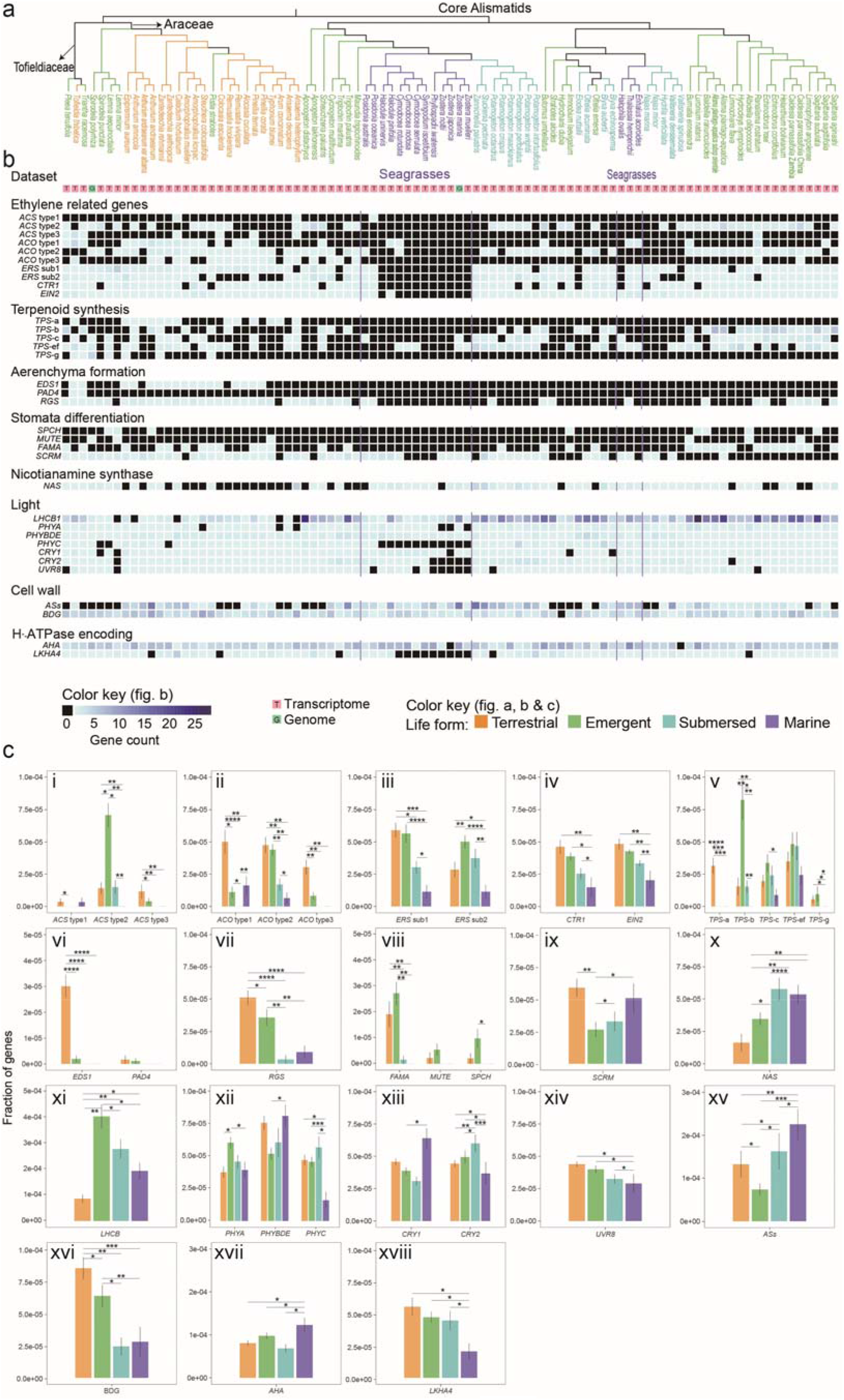
Phylogeny, gene copy counts and gene fraction of genes involved in adaptation to marine environments across Alismatales. a) Species tree of Alismatales with the relationship for the three main clades manually fixed as (core Alismatids, (Araceae, Tofieldiaceae)). b) Gene copy counts for each species. c) Gene fraction for each life-form. Gene fraction for a species was calculated by the number of corresponding genes divided by the number of total genes in a species. The gene fraction in each life-form is the average gene fraction of species in each life-form. Stars above columns represent the number of the four statistical tests (Hyperg and PhyloGLM tests based on gene presence/absence data and gene copy-number data separately) with significant differences (p-value <0.05).

Although a comprehensive phylogenetic approach was used to investigate evolutionary adaptation to the aquatic habit in this study, some genes that appear “lost” might be the product of under-expression or some other mechanisms of gene expression suppression. Further studies with more genomic data are needed to investigate these adaptations.

### KO terms related to aquatic environment adaptations

Analyses were conducted to discover KO terms for gene functions, not previously reported to be involved in the adaptation to aquatic environments. We investigated KOs that significantly enriched or depleted in the emergent, submerged and marine life-form separately (fig. 6). Most of the results are concordant with the gene evolution analyses. For example, KO terms involved in nicotianamine synthesis were enriched in the submersed life-form and the marine life-form; KO terms involving ethylene reception, phytochrome, or photosystem were depleted in the submersed and marine life-form. Moreover, we identified some depleted or enriched KOs in aquatic life-forms (e.g., elongation factor 3; supplementary table S8). These KOs have not been considered in aquatic adaptation in previous studies.

**Fig. 6.**
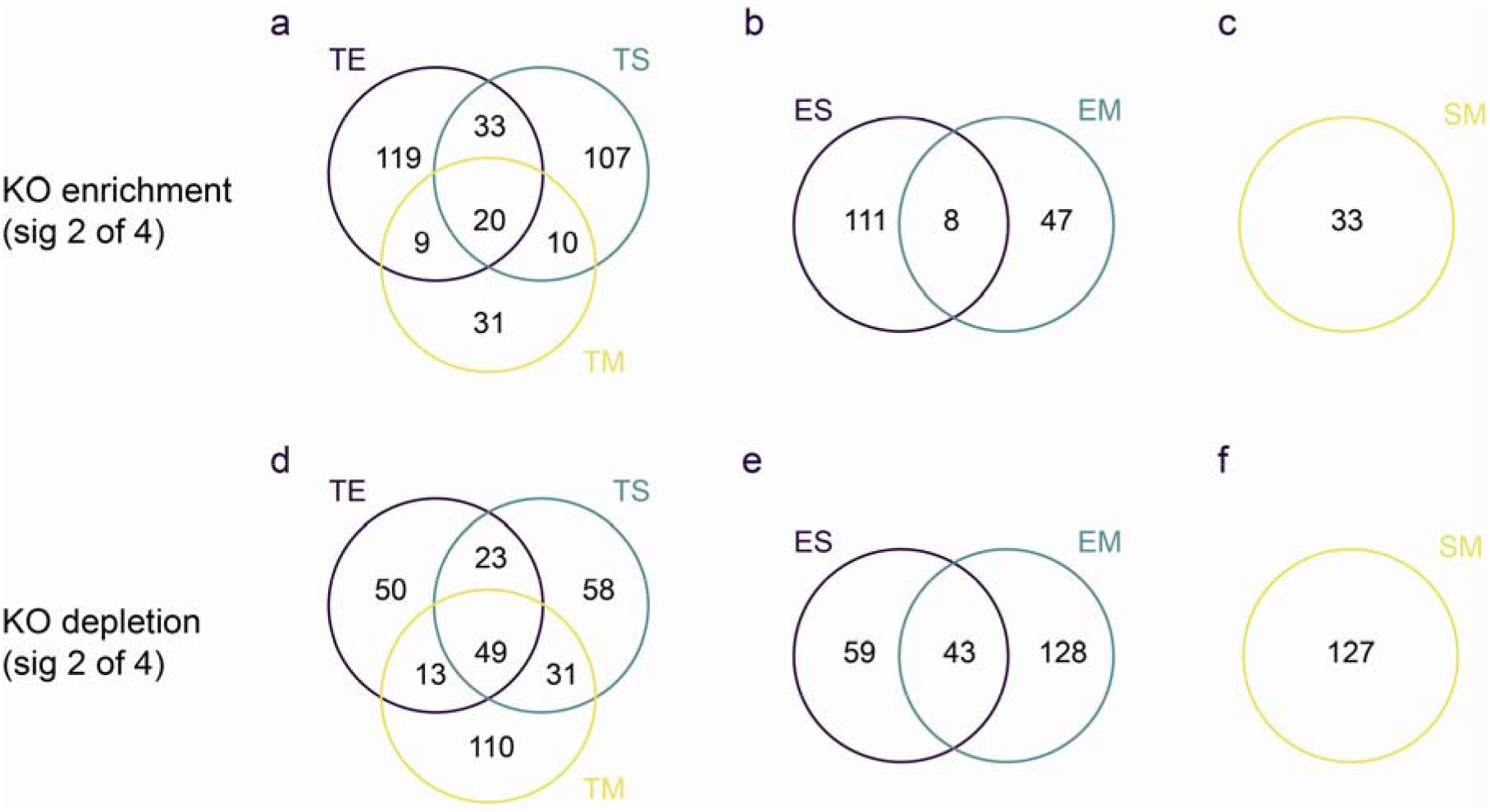
Numbers of KOs shared among life-forms. For example, in sub-figure a, the terrestrial (T) life-form was compared to the three aquatic life-forms (emergent (E), submersed (S), and marine (M)) to identify the KOs enriched in each of the three life-forms separately. Next, the number of enriched KOs shared among the three aquatic life-forms was shown in the middle of the venn. ‘Sig 2 of 4’ means two of the four statistical tests are significantly different (p-value <0.05).

Compared to the terrestrial life-form, the three aquatic life-forms shared 20 enriched KO terms (fig. 6; supplementary table S9a), which could be related to adaptation to the aquatic condition. For example, a KO term involving *CYP4X* (cytochrome P450 family 4) was enriched, although little is known about the function in plants of this KO or the others discovered. The submersed and marine life-form shared ten enriched KOs that are not shared by the emergent life-form. Among the ten KOs, one is involved in nicotianamine synthesis that plays a critical role under conditions of iron and zinc deficiency in plants (Klatte et al. 2009; Schuler et al. 2012). Another encodes the elongation factor 3, which has multiple functions in plant development and cell wall biosynthesis. Compared to the emergent life-form, the submersed and marine life-form shared eight enriched KOs (supplementary table S9b). One of the KOs is the CTF4 (chromosome transmission fidelity protein). The ortholog of CTF4 in rice sustains normal cell cycle progression (Zhang et al. 2021). The enrichment of this KO could be attributed to the well-developed aerenchyma in the submersed and marine life-forms. Compared to the freshwater submersed life-form, the marine life-form had 33 enriched KOs (supplementary table S9c). For example, three of the KOs were related to anthocyanidin metabolism and *CYP86A1* encodes the long-chain fatty acid omega-monooxygenase. The function of these KOs in plants needs further investigation to determine how they may relate to marine environment adaptation.

Compared to the terrestrial life-form, the three aquatic life-forms shared 49 depleted KOs (fig. 6; supplementary table S9d). The submersed and marine life-form shared 31 KOs that are not shared by the emergent life-form. Compared to the emergent life-form, the submersed and marine life-form shared 43 depleted KOs (supplementary table S9e). Compared to the submersed life-form, the marine life-form has 127 depleted KOs (supplementary table S9f). Five of the depleted KOs in marine life-form plants involve ethylene reception (*ETR, ERS, EIN2, EIN3*) or phytochrome, which is consistent with Olsen et al. (2016) and Lee et al. (2018). The depletion in the marine life-form also involves five photosystem KOs (*psaH, psbO, psbW, psbY*, and *psbR*), likely related to the light condition in the marine environment. The difference in light quality and intensity between submerged freshwater and marine habitats has not received much attention, in relation to gene function. The results of this research suggest differences in these habitats could have consequences for the evolution of photosystem gene function.

### Why are submersed marine angiosperms only found in Alismatales?

The angiosperms that grow entirely submersed in marine environments are all from Alismatales (Larkum et al. 2018). Here we explore possible reasons that contribute to this phenomenon: 1) The challenge of transitioning from a terrestrial to a marine environment should require a greater array of adaptations than transitioning from freshwater to seawater (Vermeij 2000). Freshwater plants have advantages over terrestrial plants regarding the adaptation to the marine environment. Although the transition from terrestrial to sea occurred in mangroves (Takayama et al. 2021), they do not grow entirely submersed, but live in coastal saline or brackish water with emergent leaves, thus reducing the need for specialized adaptations related to photosynthesis and nutrient transport. 2) Aquatic angiosperms could be divided into two categories: aquatic taxa in terrestrial orders and aquatic orders (Nymphaeales, Ceratophyllales, Acorales, and Alismatales; Du et al. 2016). During the Late Cretaceous, the Tethys seas provided vast shallow water areas (Larkum et al. 2018), which are suitable for the origination of seagrasses. On the contrary, most of the aquatic taxa in terrestrial orders originated during the Late Cenozoic and later, thus are younger than the existence of the Tethys seas, (e.g., aquatic *Ranunculus* (He et al. 2022)). Therefore, the aquatic taxa in terrestrial orders had fewer chances of adaptation to the sea. 3) Alismatales has the largest number of aquatic species and genera among all the aquatic groups, thus having more chances of adaptation to the marine environment. For example, although Ceratophyllales originated during the Cretaceous, it includes only four species and one family (The Plant List). Moreover, compared to Nymphaeales, Ceratophyllales, and Acorales, Alismatales is relatively diverse in form. For example, Alismatales have great diversity in leaf morphology (ribbon like, broad and circular, and short linear) and reproductive system (dioecy, monoecy, hermaphroditism, and dioecy and monoecy; Flora of China: http://www.efloras.org/, accessed Feb 2022). All of the seagrasses except for *Halophila* have ribbon-like leaves, which are not common among other submersed aquatic lineages. 4) The Alismatales, with its broad diversity, may have had a better chance of having the genomic predisposition for adaptation to the marine environment. For example, many of the genes hypothesized as important for marine environments can be found in submersed forms of freshwater Alismatids, such as the increase in Nicotianamine genes and *LHCB1*. The combination of functional gains and losses could lead to the specific genomic conditions for evolution to a marine environment that is then phylogenetically constrained to this lineage. Overall, a combination of these factors likely best explains why all angiosperms that grow in a submersed marine environment are all from Alismatales. Genomes of some early evolving aquatic plants such as *Ceratophyllum demersum* and *Euryale ferox* (Yang et al. 2020), and mangrove plants (Feng et al. 2021) have been recently sequenced and can be included to better elucidate the marine and freshwater adaptation.

## Materials and Methods

### Sampling and sequencing

Ninety-one Alismatales samples from 89 species (*Alisma plantago-aquatica* and *Caldesia parnassifolia* were each represented by two samples) were included. The samples represented 40 of the 57 genera (11 of the 12 families) within core Alismatids, three of the four genera within Tofieldiaceae, and 17 of the 117 genera within Araceae. The only unsampled family is Ruppiaceae, which has a consistent phylogenetic position (e.g. Li et al. 2021; Ross et al. 2016) and has a submerged life form, similar to members of the family Potamogetonaceae. Twenty-one of the Alismatales samples were terrestrial, 37 were emergent/floating aquatic, 17 were freshwater submersed, and 16 were seagrasses. Following the Angiosperm Phylogeny Group IV (The Angiosperm Phylogeny Group 2016), *Amborella trichopoda* (Amborellaceae), *Nymphaea colorata* (Nymphaeaceae), *Austrobaileya scandens* (Austrobaileyaceae) and *Acorus americanus* (Acoraceae) were included as outgroups. Of the 95 samples, four were from published whole-genome sequencing data, 32 were RNA-seq reads accessed from NCBI SRA, and 59 were RNA-seq reads (2 × 150 bp) generated in this study (supplementary table S1).

### Transcriptomic data processing and assembly

Raw read processing and organelle and over-represented reads were removed following Morales-Briones et al. (2021). After assembly with Trinity v2.8.5 (Haas et al. 2013), low quality transcripts and chimeric transcripts were removed before clustering into putative genes. Lastly, transcripts were translated with TransDecoder v5.5.0 (accessed from https://github.com/TransDecoder, Sep 2019) and transcript redundancy was assessed. See Supplementary Material online for detailed methods.

### Homology inference, chloroplast genes assembly and phylogenetic analyses

To infer homologs, an all-by-all BLASTN search was performed for CDSs of all 95 taxa. Then putative homolog groups were clustered using MCL v1.37 (van Dongen 2000). Orthologs were inferred using the ‘monophyletic outgroups’ (MO) method from Yang and Smith (2014). A maximum likelihood (ML) tree using the concatenated orthologs was built using RAxML v8.2.12 (Stamatakis et al. 2006; all ML trees in this study were inferred using RAxML with the GTRGAMMA model and 200 rapid BS replicates unless noted). A species tree was inferred from gene trees of orthologs with ASTRAL v5.7.3 (Zhang et al. 2018). Moreover, a species tree was built with 99 samples, which included genomic data of six Alismatales, transcriptomes, and five outgroups. This species tree was used to verify possible WGD events, accompanied by syntenic analyses (Supplementary Material online).

We accessed the cp genomes for 13 species of Alismatales from NCBI GENBANK (supplementary table S1). We also assembled cp CDSs using organelle reads from RNA-seq. Phylogenetic trees were produced using the RAxML and ASTRAL separately. See Supplementary Material online for detailed methods.

### Assessing phylogenetic conflict and phylogenetic networks

To investigate phylogenetic conflict, each of the nuclear gene trees was rooted using *Amborella trichopoda*. Then, each of the rooted trees with bootstrap (BS) support ≥50% for the corresponding node was mapped against the ASTRAL tree using PhyParts (Smith et al. 2015) to calculate the concordant/conflicting bipartitions and the internode certainty all (ICA) values. To distinguish the nodes lacking support from conflicting support, Quartet Sampling (QS, Pease et al. 2018) was carried out with the ASTRAL tree and the concatenated alignment of all the nuclear orthologs.

To infer species networks that account for both reticulation (e.g., hybridization) and ILS, Bayesian inference was carried out with PhyloNet for five groups, viz., Alismatales (includes 10 species), Alismataceae (14), Hydrocharitaceae (17), Potamogetonaceae (11), and Araceae (19). See Supplementary Material online for detailed methods.

### Phylogenetic hypothesis testing for Alismatales

Phylogenetic analyses detected strong conflict for the relationship between core Alismatids, Araceae, and Tofieldiaceae. To test which topology is the most favored, we carried out approximately unbiased (AU) tests (Shimodaira 2002). The 798 gene trees from PhyloNet analyses of Alismatales were used. Site-wise log-likelihood scores for each of three alternative topologies of Alismatales were calculated using RAxML. Then, AU tests were carried out using CONSEL (Shimodaira and Hasegawa 2001). As complementary analyses to the AU tests, we counted the number of genes supporting each of the three topologies using likelihood scores from RAxML. We also counted the number of genes supporting each of the three topologies using raw BS values and the BS values ≥75% in gene trees separately.

To test if the discordance among the three main clades can be explained by polytomies instead of dichotomies, we carried out the quartet-frequency-based polytomy tests by Sayyari and Mirarab (2018) as implemented in ASTRAL. We filtered the 1,005 orthologs to include only orthologs that were present in all nine Alismatales species and two outgroups. In this way, 602 orthologs were extracted. ML gene trees were inferred using RAxML separately. Then, the polytomy test was performed using gene trees with raw BS values. Because this test is sensitive to gene tree error, we performed a second polytomy test using gene trees whose branches with BS values ≤75% were collapsed.

### Divergence time estimation

Divergence time was estimated by BEAST v2.5.2 (Bouckaert et al. 2014). The topology of Alismatales was fixed based on the ML tree inferred from the 1,005-nuclear ortholog matrix. Eight fossil calibration points and one secondary point were applied (supplementary table S2). The top 40 clock-like orthologs were extracted from the nuclear orthologs and used for divergence time estimation. See Supplementary Material online for detailed information on divergence time estimation.

### Inference of genome duplication

To investigate potential WGD events, we applied tree-based and *Ks* methods. 1) Putative WGDs were assessed by counting the proportion of duplicated genes at nodes. First, ML trees were constructed for the homologs inferred from MCL. SH-like support was assessed using the SH-like test (Anisimova et al. 2011) with RAxML. Then 3,492 gene clusters including ≥30 taxa and SH-like support >80 were extracted. Last, gene duplication events were recorded by mapping the rooted gene clusters to the ASTRAL tree of Alismatales. These analyses were done using scripts from Yang et al. (2018). Putative WGDs were also detected using Tree2GD v1.0.37 (accessed from https://github.com/Dee-chen/Tree2gd in Dec. 2020). To reduce computing time, 48 samples were selected to keep the main clades from the tree of the full data set and used for the Tree2GD analysis. 2) Peptide (PEP) and CDS from paralogous pairs were used to plot within-species *Ks* distribution using the pipeline in Yang et al. (2015). *Ks* <0.02 were filtered. According to preliminary analyses, most Alismatales species have a peak at *Ks* ≈0.05 or 0.1. We ignored this peak for most species, as it is difficult to distinguish between a recent WGD and a recent retrotransposition event (Tiley et al. 2018). To determine the position of a *Ks* peak, between-species *Ks* were plotted using the pipeline in Wang et al. (2019) with minor changes. The *Ks* for nine backbone nodes of Alismatales were estimated using multiple species pairs in each node (supplementary table S5). A *Ks* plot showed that the stem node of Alismatales corresponded to a peak of *Ks* ≈1.45 (supplementary fig. S8-1a). Hence, we ignored the peaks earlier than 1.45 for all the species, as they may correspond to WGDs earlier than Alismatales. At last, three candidate WGD events were verified using syntenic genes in genome (see Supplementary Material online for detailed methods).

### Identification of KOs and genes related to adaptation to aquatic environments

We prepared two data matrices for these analyses. 1) KO annotation for PEPs/genes of all the 91 Alismatales samples was obtained using eggNOG-Mapper v2 (Cantalapiedra et al. 2021). Each query was searched against eggNOG sequences using diamond (*-m diamond*) to fix the taxonomic scope in plants (*-- tax_scope 33090*). A total of 1,718,373 (70.12 %) genes out of 2,450,555 predicted proteins were successfully annotated using the KEGG Orthology (KO) database (Kanehisa et al. 2016). Then, PEPs/genes were clustered according to the KO annotation. Last, a matrix including gene copy number per KO term was generated. 2) According to Olsen et al. (2016) and Han et al. (2019), seven groups of genes involved in ethylene reception and reaction, terpenoid synthesis, arerenchyma formation, stomata differentiation, light, cell wall, and H^+^-ATPase were selected to examine the evolution in Alismatales (supplementary table S6). Leukotriene A4 hydrolase (LTA4H) and nicotianamine synthase (NS), selected from our KO analyses, were also tested. A database consisting of PEPs of the 95 samples and a few other species, such as *Arabidopsis*, was generated. For each of these genes, at least one sequence was used as a ‘bait’ to search against the database with SWIPE v2.1.0 (Rognes 2011). A maximum of 12 hits were kept for each species. Candidate genes were filtered by using PFAM domains or InterProScan domains. After that, the candidate genes were aligned using MAFFT v7.407 (Katoh et al. 2013), and low occupancy columns were trimmed using Phyutility v.2.7.1 (-clean 0.01). An ML tree for each of these genes was built using RAxML. Subdivisions for each gene family were assigned according to positions of outgroups and previous studies. Finally, a Hypergeometric test (Hyperg) and PhyloGLM test (Levy et al. 2018) were applied to check if a KO or gene family is significantly associated with the life-form. Two versions of Hyperg and PhyloGLM tests were used, based on either gene presence/absence data (hypergbin and phyloglmbin) or gene copy-number data (hypergcn and phyloglmcn) (Levy et al. 2018). Hyperg ignores the phylogenetic structure of a data set, while PhyloGLM takes it into account (Levy et al. 2018).

## Conclusion

This study investigated the source of phylogenetic conflict within Alismatales, and adaptations to aquatic environments. The species network search and hypothesis testing recovered a relationship: ((Tofieldiaceae, Araceae), core Alismatids). Phylogenetic conflict among the three main clades could be attributed to introgression and incomplete lineage sorting. Our analyses also suggested that 18 candidate WGD events could have occurred in Alismatales, including one at the MRCA of core Alismatids. We found that common strategies in freshwater submersed species and seagrasses included loss of genes involved in terpenoid synthesis and stomatal differentiation (*SMF* subfamily) and expansion of genes involved in nicotianamine and sulfated polysaccharides syntheses. Another adaptive strategy is the expansion of H^+^- ATPases genes in seagrasses. Lineage-specific strategies included the loss of light-related genes (harvest, sense, and UV resistance) in Zosteraceae that were retained in other seagrasses. We also identified several KOs that might be related to adaptation to aquatic environments.

The molecular mechanisms for adaptation to novel environments, such as freshwater and marine aquatic habitats, are complex. This study sheds light on the potential genomic mechanism involved in the evolution of these early monocots and adaptation to aquatic environments. While a wide sampling of aquatic lineages and habits here provides means to assess evolutionary patterns within Alismatales, a more complete sampling of Araceae, in particular, can better assess the phylogenetic relationship within Alismatales. Moreover, future studies investigating aquatic adaptation could benefit from sampling a wider array of plant groups with multiple independent origins of aquatic taxa.

## Supplementary Material

Supplementary data are available at Molecular Biology and Evolution online.

High-resolution plant photos, assembled transcriptomes, sequence alignments, phylogenetic trees, and KO terms are available at the figshare: https://doi.org/10.6084/m9.figshare.16967767.

## Acknowledgments

This work was supported by the National Natural Science Foundation of China (grant numbers. 31670226, 32070231) and the Strategic Priority Research Program of Chinese Academy of Sciences, China (grant number XDB31000000), and the Sino-Africa Joint Research Center. We appreciate Duoyuan Chen, Ning Wang, and Tao Shi for suggestions on WGD analyses; Susanne S. Renner, Kuo Liao, Zhi-Zhong Li, Yu Cao, and Zhong-Ming Ye for plant sampling; Samuli Lehtonen for plant identification; Tao Wan and Can Dai for comments on this manuscript; the three anonymous reviewers for their constructive suggestions.

## Author Contributions

L.Y.C., J.M.C., and Q.F.W designed research; L.Y.C., M.L.M., F.L., G.G.H., J.M.C., and Q.F.W. contributed to the taxon sampling and sequencing; L.Y.C., D.F.M-B., and J.X.H. performed DNA/RNA data processing, phylogenetic analyses, WGD analyses and molecular dating; B.L. and L.Y.C. performed analyses on adaptation to aquatic environments; L.Y.C., D.F.M-B., and B.L. prepared the figures and tables. L.Y.C. drafted the manuscript. All authors revised the manuscript.

