## Supplementary Fig. S3 for "Phylogenomic analyses of Alismatales shed light into adaptations to aquatic environments"

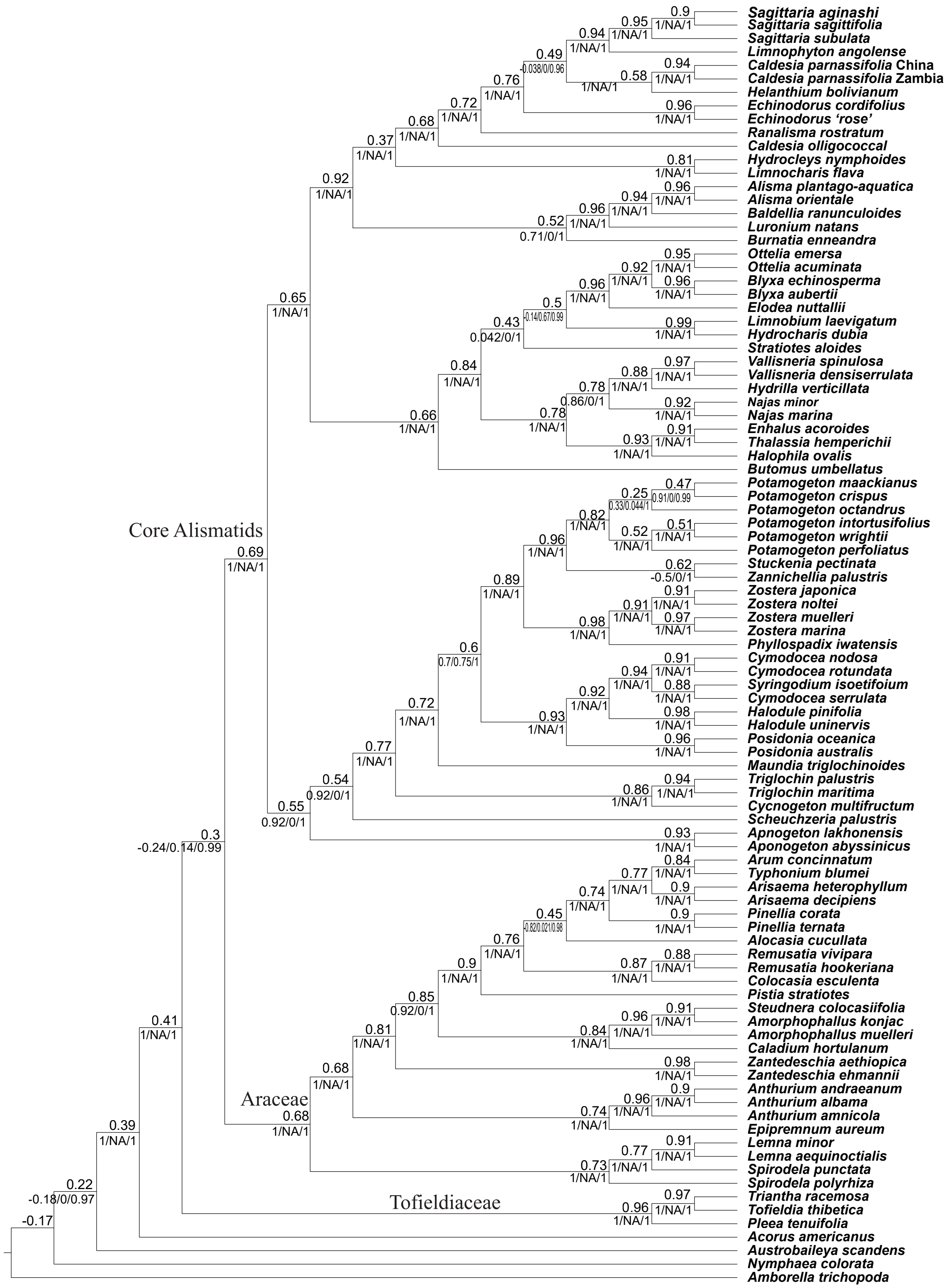


Figure S3. Species tree of Alismatales inferred from ASTRAL with 1,005 nuclear genes. Numbers above branches indicate the internode certainty (ICA) score. Numbers below branches indicate the Quartet Sampling (QS) scores (Quartet Concordances, Quartet Differential, and Quartet Informativeness).
