## Supplementary Fig. S4 for "Phylogenomic analyses of Alismatales shed light into adaptations to aquatic environments"

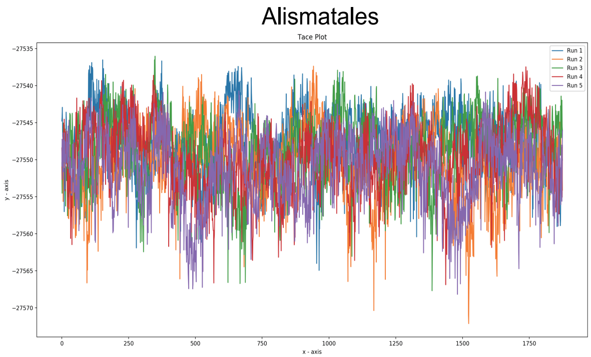


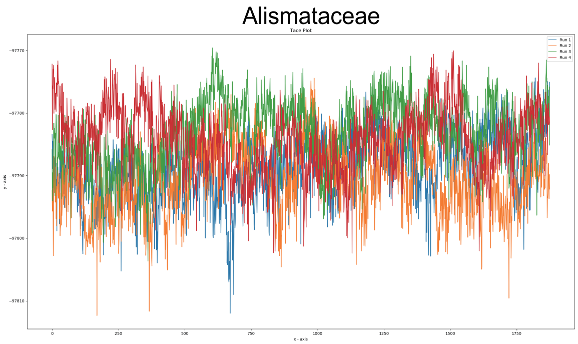


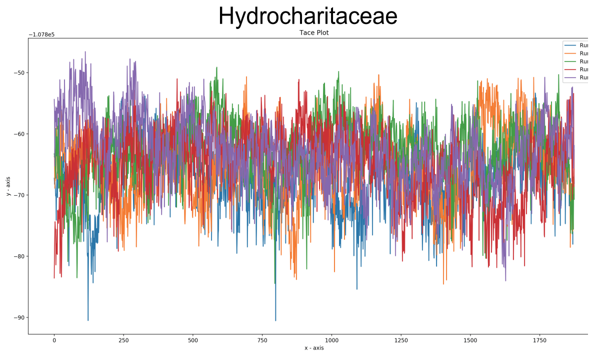


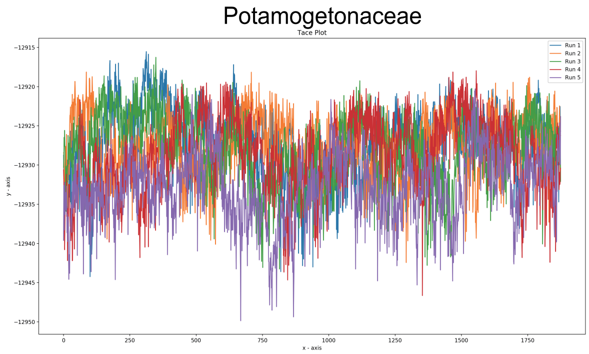


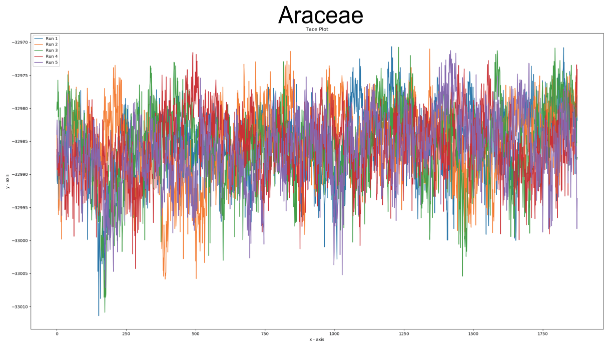


Figure S4. Convergence checking for PhyloNet analyses of the five reduced taxon data of Alismatales. MCMC chains of 20, 30, 20, 15, and 40 million with sample frequency of 2,000, 3,000, 2,000, 1,500, and 4,000 were carried out for the five groups. Five independent runs were applied for Alismatales, Hydrocharitaceae, Potamogetonaceae separately, and all the runs converged. Ten independent runs were applied for Alismataceae and Araceae separately, but only four and five runs were converged for the two families separately.
