## Supplementary Fig. S5 for "Phylogenomic analyses of Alismatales shed light into adaptations to aquatic environments"

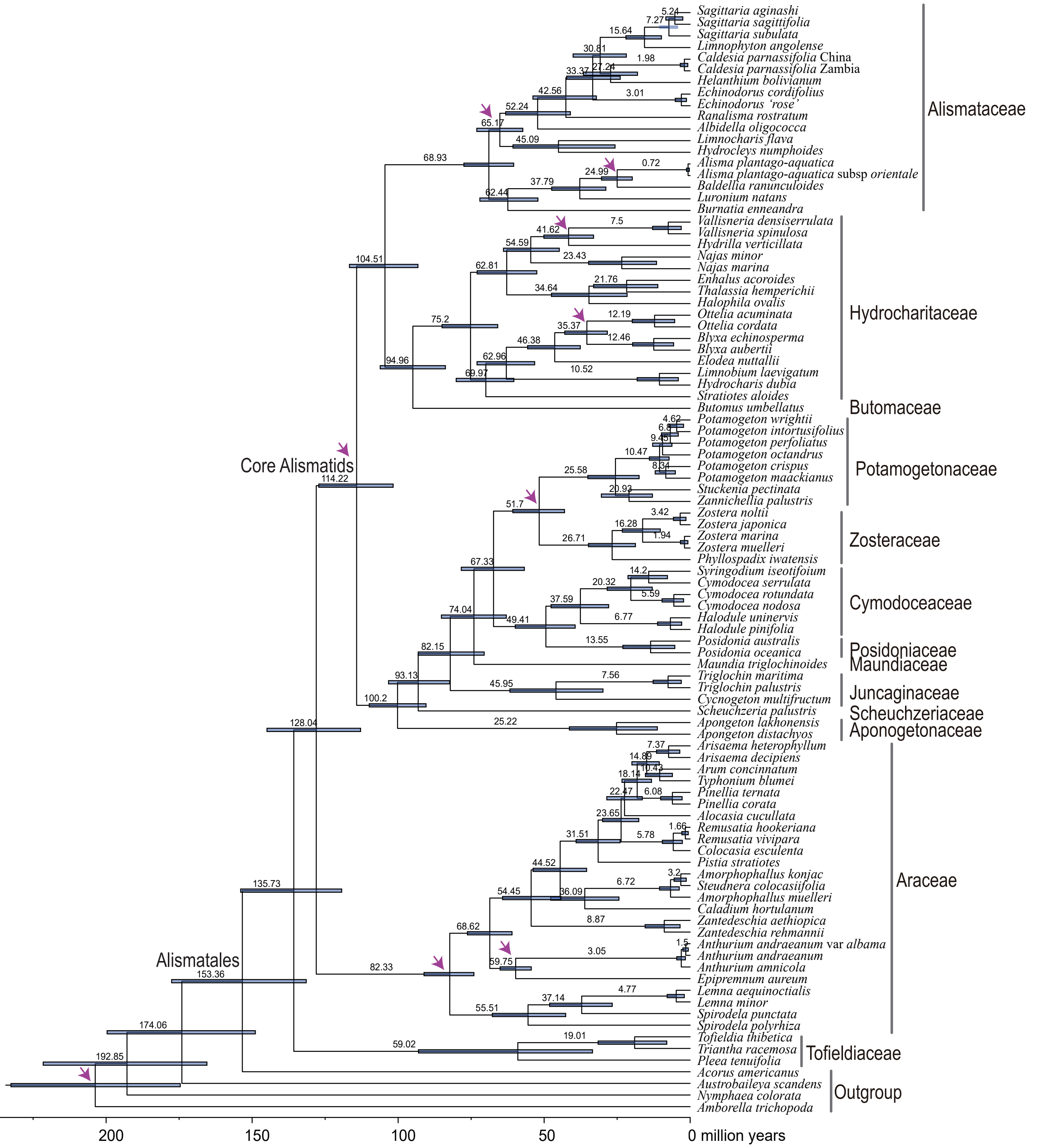


Figure S5. Chronogram of Alismatales inferred from 40 clock-like genes selected from the 1,005 MO nuclear genes. Arrows present the nine calibration points.
