## Supplementary Fig. S6 for "Phylogenomic analyses of Alismatales shed light into adaptations to aquatic environments"

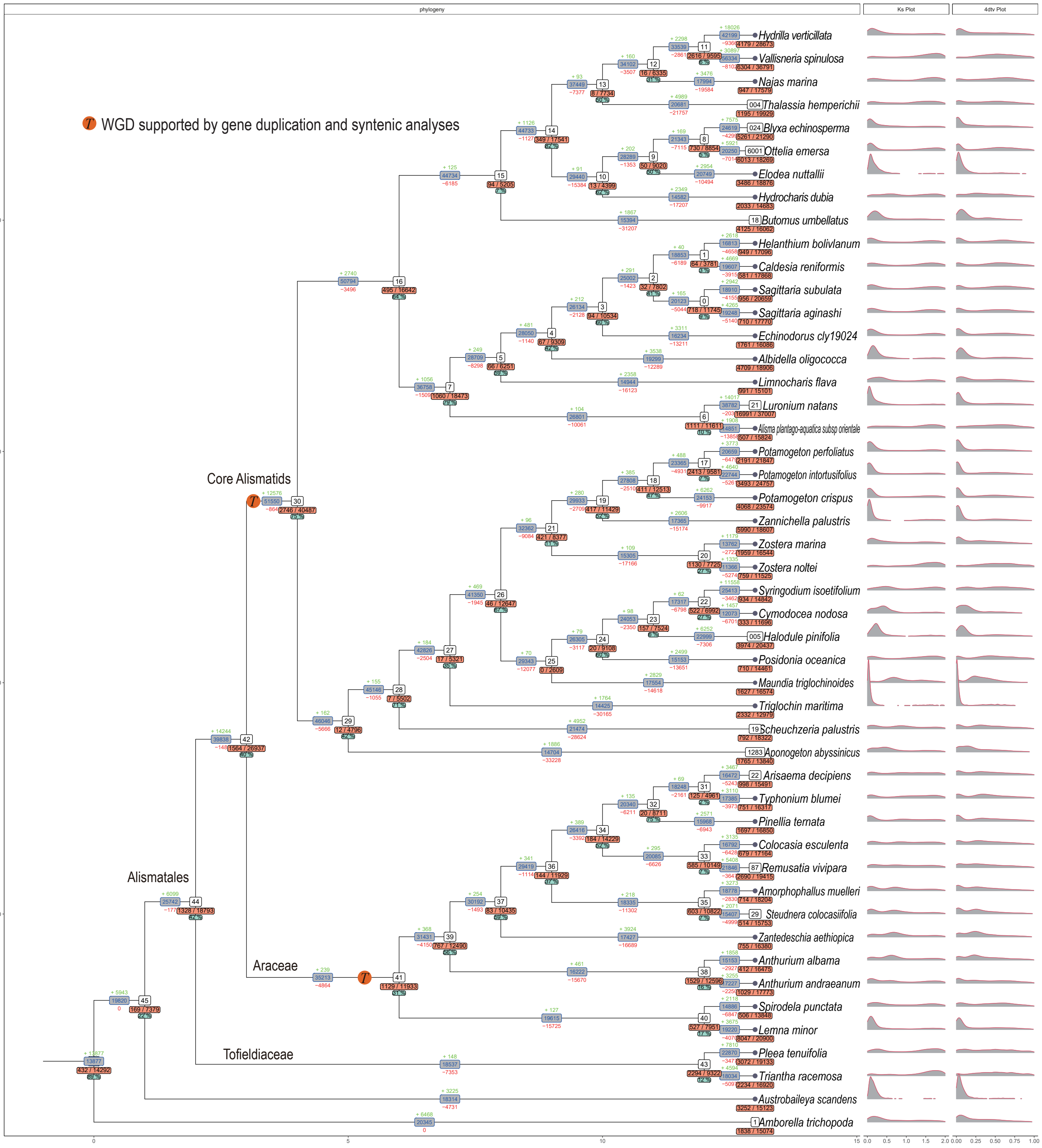


Figure S6. Genome duplication inferred from 48 species using Tree2GD. Blue numbers in grey boxes on branch lines represent the numbers of gene families descending from ancestors. Numbers above (green) and under (red) the grey boxes represent the numbers of gene family expansions and gene family contractions respectively. Numbers in white boxes represent node numbers. Numbers before and behind the virgule (/) in the orange box respectively represent the number of gene duplications shared by the taxa descending from the branch and the total gene number in an ancient clade before a WGD experiencing along the branch. The percentage in the green box represents the proportion of gene pairs kept at the node. Some of the node numbers correspond to those in Fig. 2.
