## Supplementary Fig. S7 for "Phylogenomic analyses of Alismatales shed light into adaptations to aquatic environments"

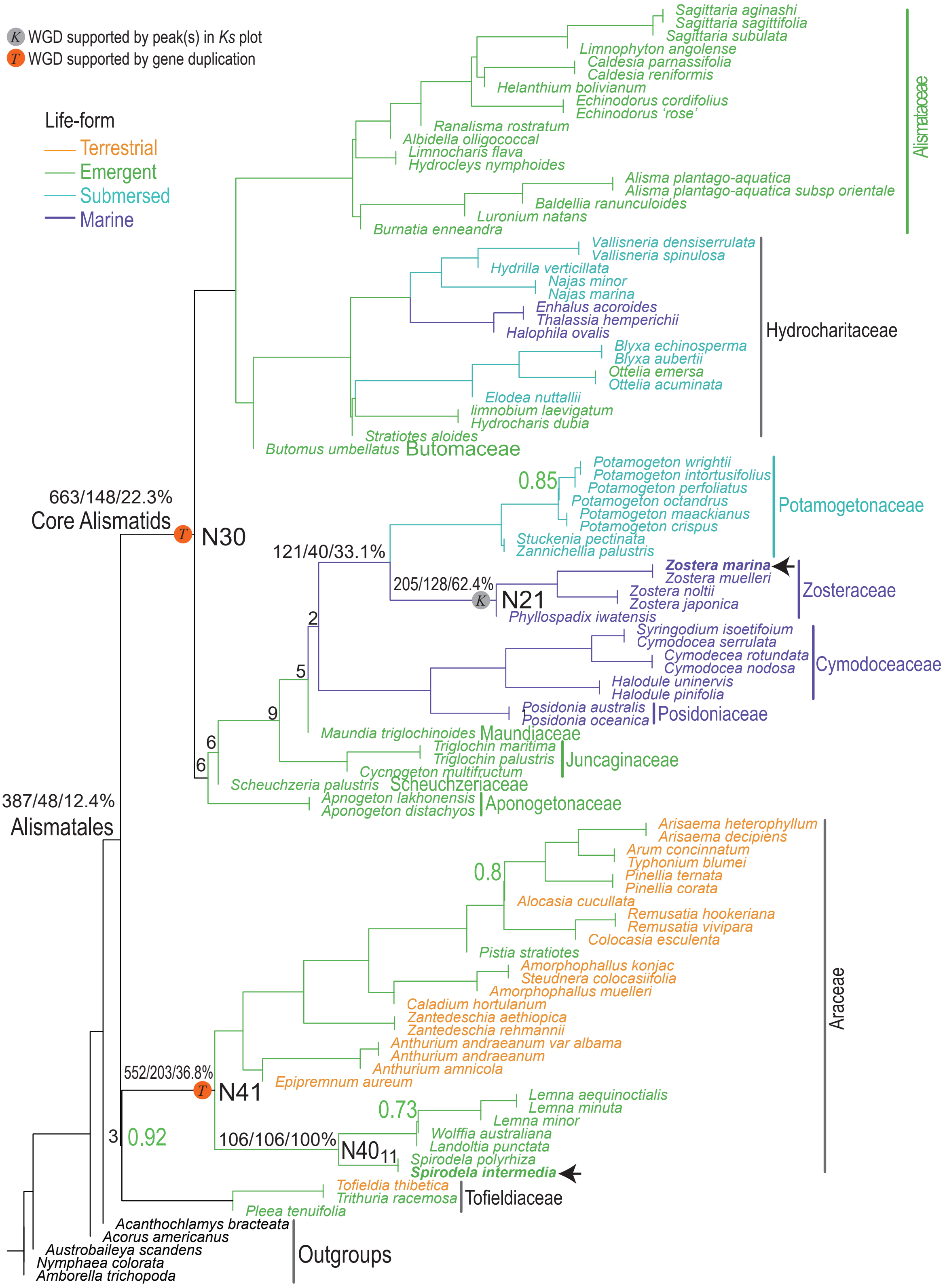


Figure S7. Verification of three WGD events using syntenic analyses and 99 samples. The ASTRAL species tree was inferred from 1,050 orthologs in 99 samples. Branches and species names were colored according to the corresponding life-forms. Node numbers correspond to those in Fig. 2. Arrows represent that syntenic analyses were carried out for the two species (*Zostera marina* and *Spirodela intermedia*). All nodes have ASTRAL local posterior probability =1 unless noted near the branches (green). 3,493 gene clusters were mapped to the ASTRAL tree using PhyParts to get duplicated clusters at each node. Number separated by the virgule(/) represents the number of duplicated clusters at the node, numbers, and proportion of clusters that have duplicated and collinear genes. For example, 663/148/22.3% at N30 represent 663 out of 3,493 clusters duplicated at the node. Among the 663 gene clusters, 148 (22.3%) have duplicated and collinear genes in *Z. marina*. When the number of duplicated clusters <100, only the number (black) was shown. For example, the node formed by Araceae and Tofieldiaceae. Collinear genes at N41 and N40 were counted using *S. intermedia*, while nodes Alismatales, N30, (Potamogetonaceae + N21), and N21 were counted using *Z. marina*.
