## Supplementary Fig. S8-2 for "Phylogenomic analyses of Alismatales shed light into adaptations to aquatic environments"

# *Ks* 0.02–3.0

### c. Hydrocharitaceae

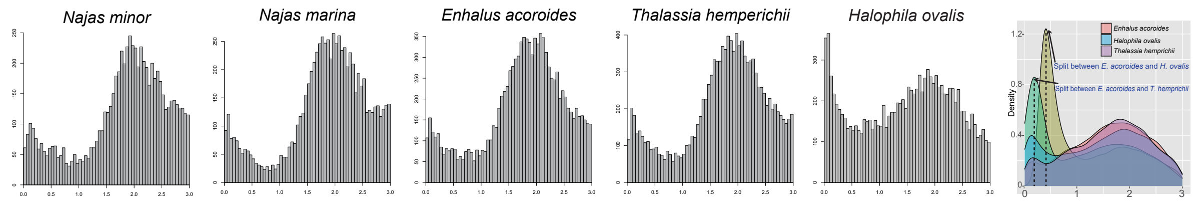


### d. Butomaceae, Potamogetonaceae, Aponogetonaeae

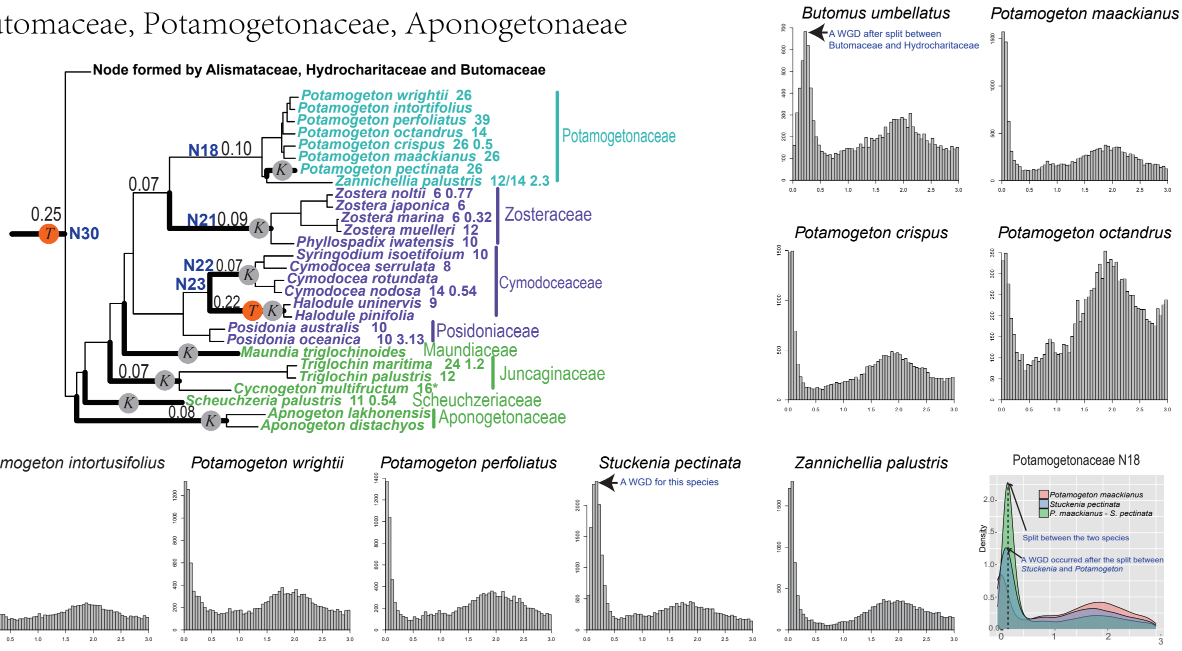


### e. Zosteraceae

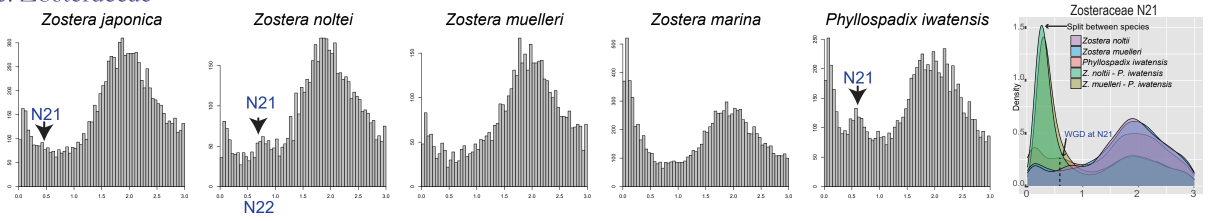


### f. Cymodoceaceae

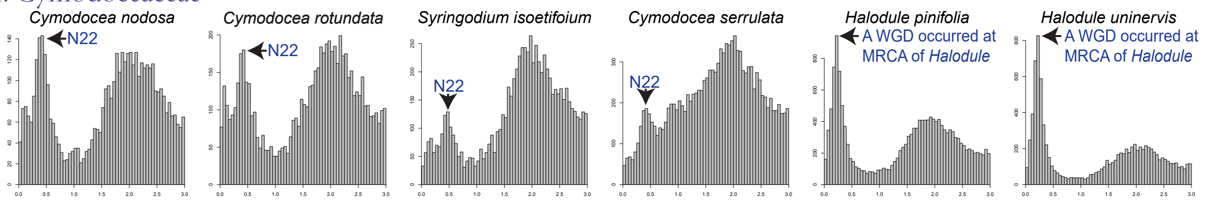


### g. Posidoniaceae and Maundiaceae

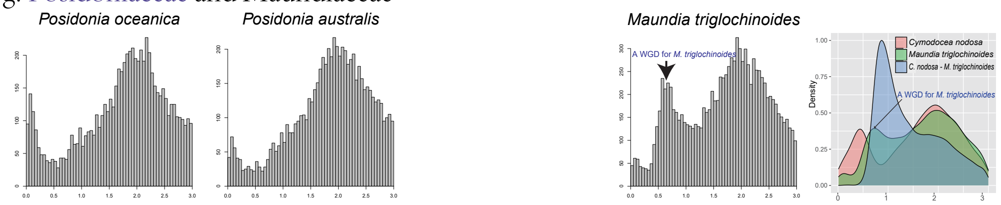


### h. Juncaginaceae

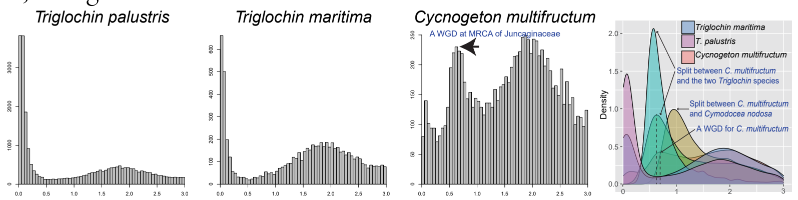


Figure S8-2. Distribution of *Ks* values range from 0.02–3.0 for families Hydrocharitaceae, Butomaceae, Potamogetonaceae, Aponogetonaeae, Zosteraceae, Cymodoceae, Posidoniaceae, Maundiaceae and Juncaginaceae.
