## Supplementary Fig. S8-3 for "Phylogenomic analyses of Alismatales shed light into adaptations to aquatic environments"

# *Ks* 0.02–3.0

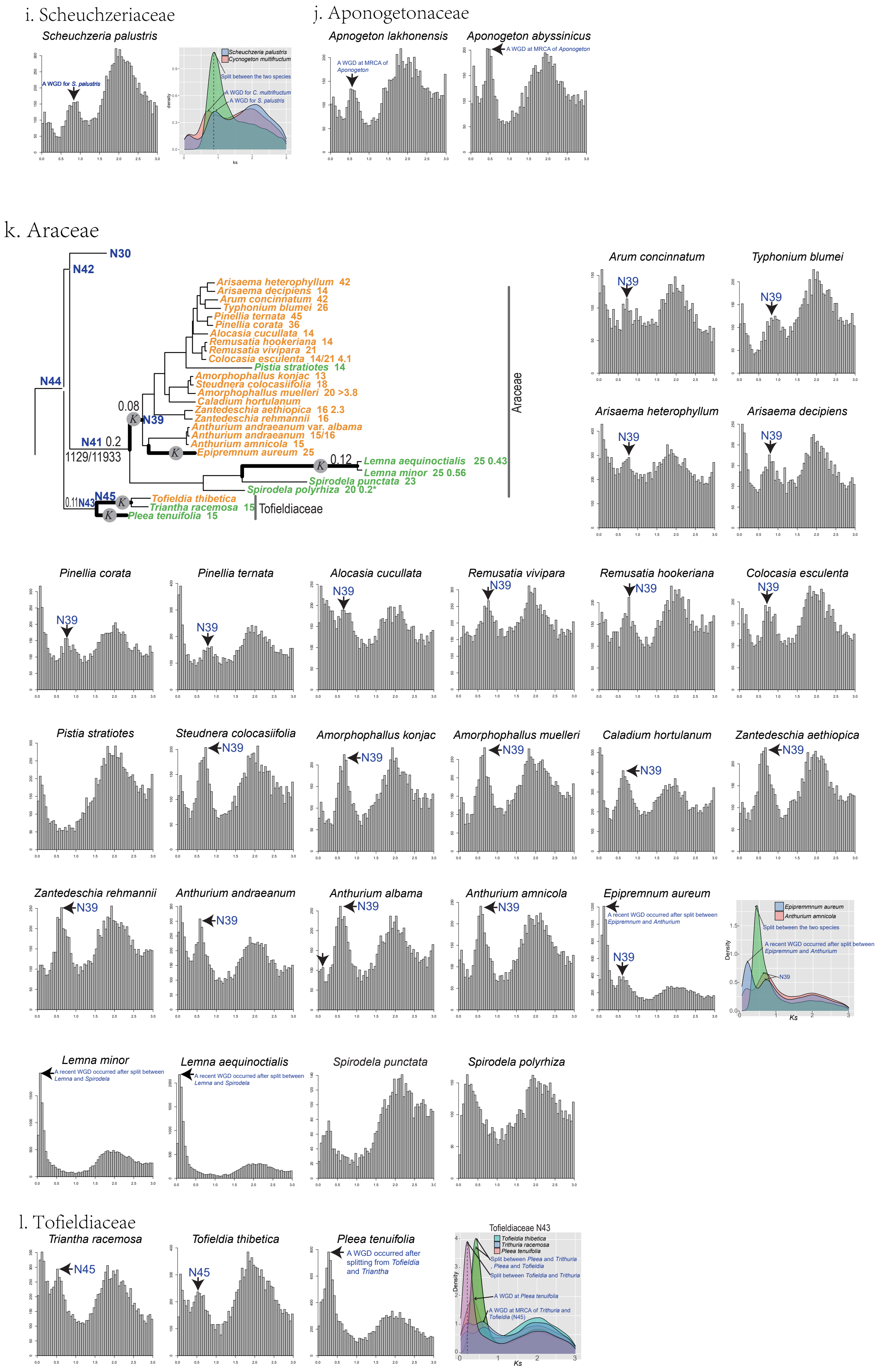


Figure S8-3. Distribution of *Ks* values range from 0.02–3.0 for families Scheuchzeriaceae, Araceae and Tofieldiaceae.
