## Supplementary Fig. S8-4 for "Phylogenomic analyses of Alismatales shed light into adaptations to aquatic environments"

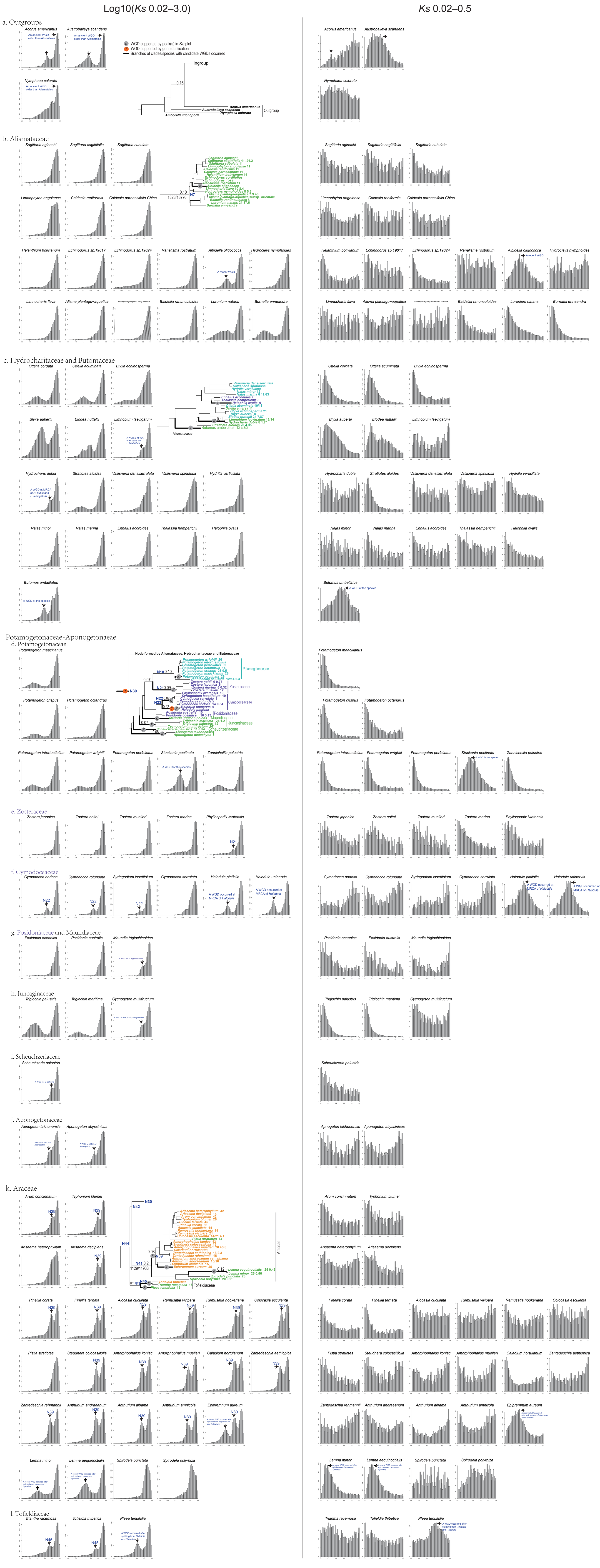


Figure S8-4. Distribution of *Ks* values with log10 transformed and range from 0.02–0.5. The white and grey plots show the within species *Ks* estimated using the pipeline in Yang et al. (2015); the color plots show the density of within species *Ks* and between species *Ks* estimated using the pipeline in Wang et al. (2018). Phylogenetic trees were cited from Fig. 2, in which decimals above branches show the proportaion of duplicated clusters at the node using the pipeline in Yang et al. (2018). Numbers below branches show the proportion of duplicated genes at the node recovered using Tree2GD results (supplementary fig. S6). Node numbers such as N43 also refer to the nodes in Fig. 2 and supplementary fig. S6. Numbers next to species names indicate chromosome number and 1c values assessed from IPCN Chromosome Reports (http://legacy.tropicos.org/Project/IPCN) and Plant DNA C-value Database (http://data.kew.org/cvalues/) respectively.
