## Supplementary Fig. S1 for "Phylogenomic analyses of Alismatales shed light into adaptations to aquatic environments"

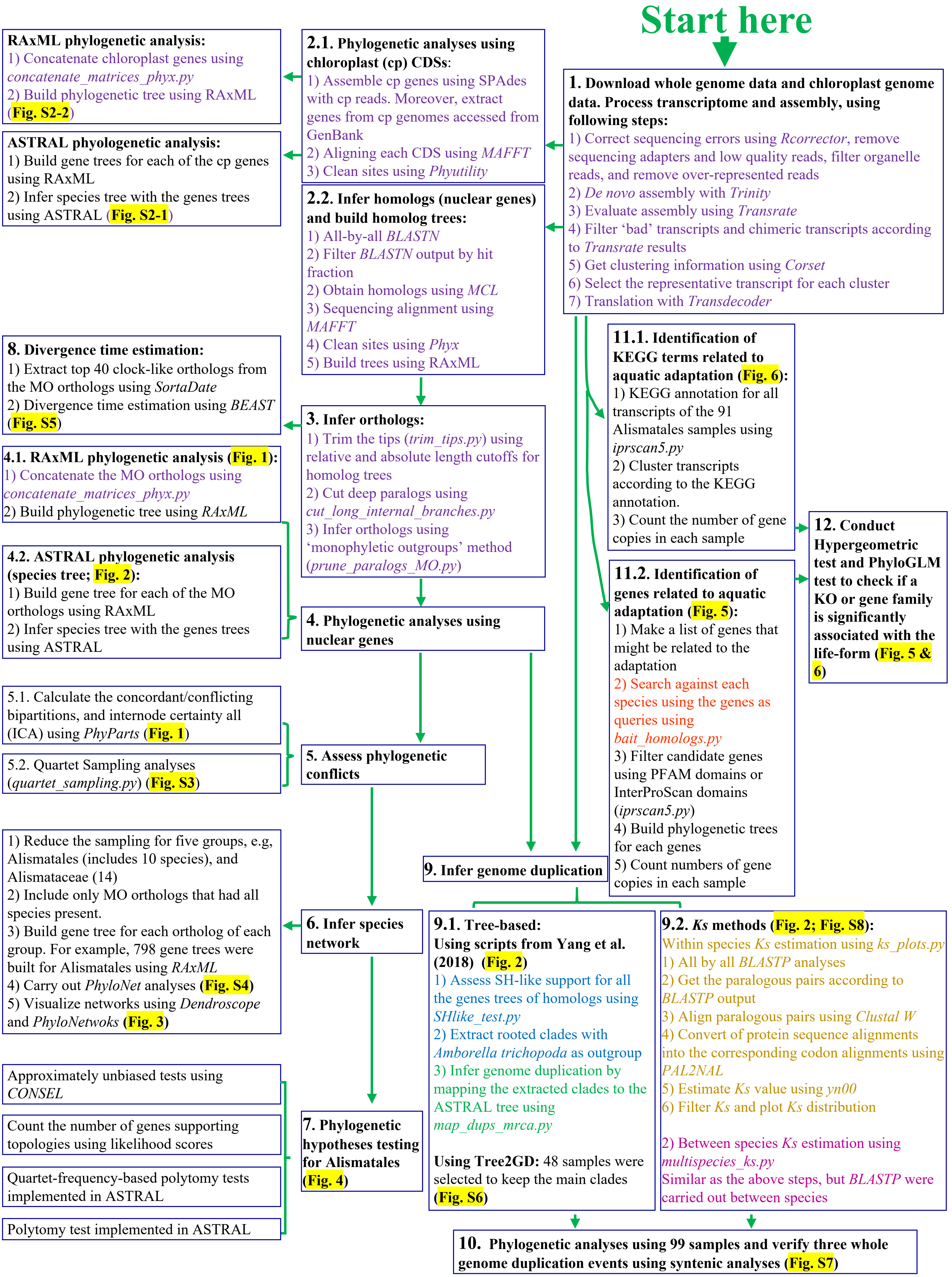


Purple indicates analyses that were assisted using scripts from https://bitbucket.org/yanglab/phylogenomic_dataset_construction/src/master/
Assisted using scripts from https://bitbucket.org/ningwang83/portulacineae/src/master/
Assisted using scripts from https://https://bitbucket.org/yangya/caryophyllales_mbe_2015/src/master/
Assisted using scripts from https://bitbucket.org/yangya/genome_walking_2016/src/master/
Assisted using scripts from https://bitbucket.org/blackrim/clustering/src/master/
Black indicate analyses that were mainly conducted according to manuals of corresponding software, and *SHELL* or *R* scripts

**Supplementary Fig. S1.** Work flow of data analyses in this study.
