## Supplementary Fig. S2-1 for "Phylogenomic analyses of Alismatales shed light into adaptations to aquatic environments"

Coalescent-based species tree (ASTRAL)

53 chloroplast CDSs

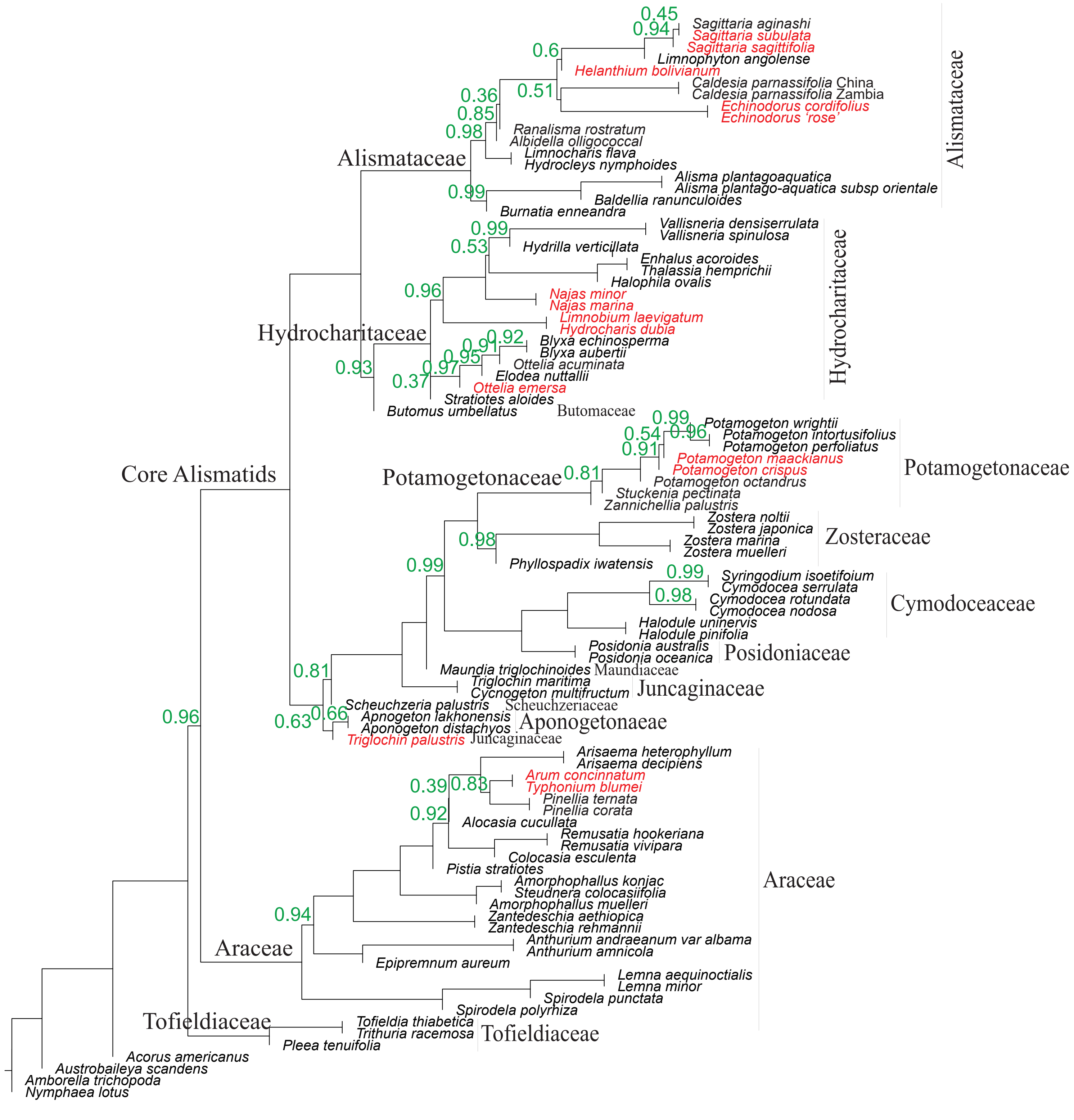


Figure S2-1. A cladogram of Alismatales inferred from ASTRAL analysis with chloroplast CDSs. Numbers (green) near branches indicate local posterior probabilities (LPP). The numbers were not shown when LPP = 1. The taxon name was red colored when its phylogenetic position/relationship is inconsistent with the ASTRAL tree inferred from 1,005 orthologs (fig. 2).
