## Supplementary Material for "Phylogenomic analyses of Alismatales shed light into adaptations to aquatic environments"

**Transcriptomic data processing and assembly**

Raw reads processing, assembly, and translation followed the pipeline of Yang and Smith (2014; <https://bitbucket.org/yanglab/phylogenomic_dataset_construction/>) and Morales-Briones et al. (2021) with minor modifications. Sequencing errors in raw reads were corrected with Rcorrector v1.0.4 (Song and Florea 2015). Adapters and low-quality bases were removed with Trimmomatic v0.27 (SLIDINGWINDOW:4:15 LEADING:5 TRAILING:4 MINLEN:80 (Bolger et al. 2014). Additionally, organelle reads were filtered with Bowtie2 v2.3.5 (Langmead and Salzberg 2012) by mapping to Magnoliophyta organelle genomes accessed from the Organelle Genome Resources database (RefSeq; (Clark et al. 2007); accessed on October 17, 2018) as references. Over-represented reads were detected by FastQC v0.11.8 (https://www.bioinformatics.babraham.ac.uk/projects/fastqc/) and removed. After these processes, we recovered 4.9–31.5 million (average16.4) read pairs for each of the 59 newly generated transcriptomes and 4.9–31.5 million (average 22.6) read pairs for transcriptomes assessed from NCBI SRA (supplementary table S1). *De novo* assembly was carried out using Trinity v2.8.5 (Haas et al. 2013) with default settings. Assembly quality was accessed with Transrate v1.0.3 (Smith-Unna et al. 2016). Low quality transcripts were removed based on any of the three cut-off values, Cord ≤ 0.50, Ccov ≤ 0.25 and Cnuc ≤ 0.25. Chimeric transcripts were removed following the methods in Yang and Smith (2013) by using genomes of *Zostera marina*, *Spirodela polyrhiza* and *Oryza sativa* as references (Yang and Smith 2013). Filtered reads were mapped to filtered transcripts using Salmon v0.9.1 (Patro et al. 2017), and putative genes were clustered using Corset v1.09 (Davidson and Oshlack 2014). Then the longest transcript within each putative gene was retained according to our previous study (Chen et al. 2019). To find open reading frames (ORFs), retained transcripts were translated with TransDecoder v5.5.0 (accessed from <https://github.com/TransDecoder>, Sep 2019) by using *Z. marina*, *S. polyrhiza* and *Oryza sativa* as references. Redundant ORFs were reduced with CD-HIT v4.8.1 (-c 0.99; Li and Godzik (2006)).

**Homology and orthology inference and phylogenetic analyses**

Homology inference was carried out following Yang and Smith (2014) and Morales-Briones et al. (2021) with minor modifications. First, an all-by-all BLASTN search was performed for the CDS of all 95 samples with settings ‘-evalue 10 -max_target_seqs 1000’. BLAST output was filtered with a hit fraction of 0.25. Then, putative homolog groups were clustered using MCL v14-137 (van Dongen 2000) with a log-transformed E-value cutoff of 5 and an inflation value of 1.4. Clusters with at least 70 species were retained. Each cluster was aligned by using MAFFT v7.407 (Katoh and Standley 2013) with settings ‘--genafpair --maxiterate 1000’ (sequences in this study were aligned using MAFFT with these parameters unless noted). Aligned columns with more than 90% missing data were removed using Phyx (Brown et al. 2017). The ML tree for each alignment was built. Spurious tips were trimmed by using relative (value 8) and absolute length cutoffs (value 0.4). An internal branch longer than 0.5 and a minimum number of 82 taxa were used to cut the deep paralogs. Orthology inference was carried out separately using the ‘monophyletic outgroup’ (MO, Yang and Smith 2014). The MO method generated 1,005 nuclear genes.

Concatenation and coalescent methods were used for phylogenetic analyses. Each orthologous group was aligned with MAFFT, then trimmed with Phyx. The matrix of the concatenated MO genes included 2,046,379 aligned columns with a gene and character occupancy of 93% and 79%, respectively. ML inference for the concatenated matrix was carried out using RAxML, partitioned by gene or not, using the concatenated matrix from the MO method (all ML trees in this study were inferred using RAxML unless noted). To obtain species trees, ML inference was carried out for individual alignments separately. Then, a species tree was produced from the ML trees of MO orthologs with ASTRAL v5.7.3 (Zhang et al. 2018).

**Chloroplast genes assembly and phylogenetic analyses**

We accessed the whole chloroplast genomes for 13 species from NCBI GENBANK (supplementary table S1). Seventy-seven chloroplast genes, which existed in all of these species, were extracted. Moreover, we assembled chloroplast genes. The organelle reads for each species from the filtering step were mapped to each of the 77 genes with Bowtie2 v2.3.5. Then, the chloroplast genes for each species were assembled using the mapped reads with SPAdes v3.13.1 (Bankevich et al. 2012). The assembled contigs were orientated by comparing to the reference genes using Geneious v11.1.5 (Kearse et al. 2012) and the contig with the highest similarity to other species was retained when multiple contigs were assembled for a gene. Twenty-four genes from the 77 genes were removed due to low sequence occupancy. Moreover, *Luronium natans*, *Caladium hortulanum* and *Anthurium andraeanum* were removed, as their phylogenetic position is obviously different from that in previous studies in our preliminary analyses, which implied contaminations, misassembly, or RNA-editing.

Each gene was aligned using MAFFT. Each alignment was trimmed with Phyutility, then visually inspected in Geneious. Alignments were concatenated. The chloroplast data set included 53 genes from 92 samples. The concatenated matrix included 55,716 aligned columns with a gene and character occupancy of 78% and 68% respectively. A maximum likelihood (ML) tree of the concatenated matrix was built by using the alignment partitioned by gene and 200 rapid bootstrap value (BS) replicates.

**PhyloNet analyses**

Given computational restrictions and our focus on clades that show a clear signal of conflict (i.e., low ICA and QS scores), we reduced our sampling to five groups, viz., Alismatales (includes 10 species), Alismataceae (14), Hydrocharitaceae (17), Potamogetonaceae (11), and Araceae (19). We included only orthologs that had all species present. In this way, we included 798, 694, 568, 459, and 437 orthologs for the five groups respectively. Searches were carried out allowing up to three reticulation events. MCMC chains of 20, 30, 20, 15, and 40 million with sample frequency of 2,000, 3,000, 2,000, 1,500, and 4,000 were carried out for the five groups respectively. Five independent runs were applied for Alismatales, Hydrocharitaceae, and Potamogetonaceae separately. Ten independent runs were applied for Alismataceae and Araceae separately. The first 25% of the iterations were set as burn-in. Searches were done using a cold chain with a temperature 1.0 and two hot chains with temperatures 2.0 and 3.0, respectively. Pseudo-likelihood was applied to speed up the searches. PhyloNet calculated inheritance probabilities (γ) that represent the proportion of genes contributed by each parental population to a given hybrid node (Solís-Lemus and Ané. 2016). Networks were displayed using Dendroscope v3 (Huson et al. 2012) and PhyloNetwoks (Solís-Lemus et al. 2017).

**Divergence time estimation**

The size of phylogenomic datasets makes divergence-time estimation with the entire dataset intractable, and underlying topological and rate heterogeneity across genes makes model misspecification a real concern (Smith et al. 2018)). Using SortaDate ((Smith et al. 2018); assessed in Jan 2021), we estimated the tip-to-root variation and bipartition support for gene trees of the 1,005 MO orthologs. We used the root-to-tip variance first, then tree length, and then bipartition (command ‘--order 1,2, 3’). Forty clock-like orthologs were selected and used for divergence time estimation.

Divergence time was estimated by BEAST v2.5.2 (Bouckaert et al. 2014). Eight fossil calibration points were applied and one point (supplementary table S2) was applied based on the crown age of angiosperms (Li et al. 2019). Calibrations were only applied to the nodes with high support in phylogenetic analyses. The Gamma site model was applied with estimated Substitution Rate, estimated Proportion Invariant and Subst model, Relaxed clock log-normal model, and log-normal priors. Analyses were run for one billion generations and sampled every 10,000 generations. During the MCMC chain, the tree was fixed with the ML tree inferred from the MO orthologs. The effective sample size (ESS) scores for all relevant estimated parameters were checked to ensure values above 200 using Tracer v1.7.1 (Drummond and Rambaut 2007). The first 10% of trees were discarded as burn-in, and the remaining trees were used to generate a summary tree with TreeAnnotator v2.6.3 (Drummond and Rambaut 2007).

**Phylogenetic analyses using 99 samples and verify whole genome duplication using syntenic analyses**

We started data analyses of this study at the end of 2019. At that time, genomic data for many species such as *Wolffia australiana* has not yet available. Therefore, genomic data of only two species within Alismatales were used for most analyses in this study. Up to Feb 2022, genomic data for six species in Alismatales have been available. As complementary analyses, we made a dataset with 99 samples, which include genomic data of six Alismatales species, transcriptomic data of 88 Alismatales species, and five outgroups (supplementary Table S1). Then, we identified orthologs using the all-by-all BLASTN search and the MO method and built an ASTRAL species tree as described above.

To verify WGD event occurred at the MRCA of core Alismatids (N30), MRCA of Zosteraceae (N21), and MRCA of Araceae (N41), syntenic analyses were carried out. We first screened duplicated clusters at each node of Alismatales by mapping 3,493 gene trees to the species tree using PhyParts v. 0.0.1 (Smith et al. 2015). Then, we carried out syntenic analyses to identify collinear genes in *S. intermedia* and *Z. marina* using MXScan (Wang et al. 2012). Collinear genes in *Wolffia australiana*, *Spirodela polyrhiza*, *L. minor* and *L. minuta* were not investigated as chromosome-level genomes were unavailable (we emailed authors of these genomes, but no response was obtained). Last, we examined the proportion of gene pairs in duplicate gene clusters that are collinear genes.
